# Cell cycle-dependent organization of a bacterial centromere through multi-layered regulation of the ParABS system

**DOI:** 10.1101/2023.06.27.546657

**Authors:** Jovana Kaljević, Tung B. K. Le, Géraldine Laloux

## Abstract

The accurate distribution of genetic material is crucial for all organisms. In most bacteria, chromosome segregation is achieved by the ParABS system, in which the ParB-bound centromere *parS* is actively partitioned by ParA. While this system is highly conserved, its adaptation in organisms with unique lifestyles and its regulation between developmental stages remain unexplored. *Bdellovibrio bacteriovorus* is a predatory bacterium proliferating through polyploid replication and non-binary division inside other bacteria. Our study reveals the subcellular dynamics and multi-layered regulation of the ParABS system, coupled to the cell cycle of *B. bacteriovorus*. We found that ParA:ParB ratios fluctuate between predation stages, their balance being critical for cell cycle progression. Moreover, the *parS* chromosomal context in non-replicative cells, combined with ParB depletion at cell division, critically contribute to the unique cell cycle-dependent organization of the centromere in this bacterium, highlighting new levels of complexity in chromosome segregation and cell cycle control.

**Author summary:** The precise distribution of genetic material to the progeny is essential for all living organisms, and the ParABS system is critical for this process in bacteria. Our study provides novel insights into the regulation of this system in a predatory bacterium that exhibits a non-canonical cell cycle, in which the chromosome is copied and segregated multiple times during growth inside a prey bacterium. Our work reveals the subcellular dynamics and multi-level regulation of the ParABS system in this bacterium, which results in the unique cell-cycle dependent assembly of the segregation complex at the chromosomal centromere. Our findings provide a deeper understanding of the adaptation of the ParABS system across bacterial species and developmental stages.

## Introduction

The precise segregation of genetic material during cellular division is a fundamental across all domains of life. In eukaryotes, sister chromosomes are tethered to the spindle during mitosis via kinetochores, which are protein complexes that assemble at the chromosomal centromeres. The kinetochores serve as attachment points for the spindle to pull chromosomes apart. Bacteria exploit a conceptually analogous strategy to partition copies of their chromosomal DNA into future daughter cells [1].

In most bacterial lineages, chromosome segregation is achieved by the active partitioning of the duplicated chromosomal origins (*oriC*) via the ParABS system [2,3]. The leading player in this system, the DNA-binding CTPase ParB, loads onto the chromosome by binding to *parS* sites - short palindromic centromere-like sequences, usually found near the *oriC* [2,4–8]. Nucleation of ParB-CTP on *parS* allows ParB to spread away and cover adjacent DNA, leading to the accumulation of ParB at the centromere and the formation of a higher-order nucleoprotein structure known as the partitioning complex [9–14], reminiscent of the eukaryotic kinetochore. CTP hydrolysis favors the opening of the ParB clamp and removal from the DNA, replenishing the pool available for binding on one or more *parS* sites [11,14,15]. Upon initiation of chromosome replication at *oriC*, a second partitioning complex assembles on the duplicated centromere, which will be segregated by the ParABS system. Briefly, ParB·*parS* interacts with ParA, a protein that dimerizes in its ATP-bound form and associates non-specifically with the DNA, therefore coating the entire nucleoid [16,17]. The partitioning complex stimulates the ATPase activity of ParA^ATP^, displacing it from the *oriC*-proximal edge of the DNA-bound “cloud” and generating a ParA^ATP^ gradient towards the opposite cell pole. Repeated ParA-ParB interactions drive the movement of the centromere towards the highest ParA^ATP^ concentration and across the cell, followed by the rest of the sister chromosome [3,18–21].

In several organisms, this process is enhanced by specific landmark polar proteins that either anchor the chromosomal *oriC* [22–25] or sequester the released ParA^ADP^ monomers to maintain the ParA^ATP^ gradient [26–28]. These mechanisms play a role in ensuring the unidirectionality of chromosome segregation and establishing polarity in future daughter cells [29]. Bacteria also evolved diverse strategies to precisely couple the positioning and timing of cell constriction with the segregation of either *oriC* [30] or the chromosomal terminus (*ter*) [31,32]. Consequently, segregation of bacterial chromosomes is tightly coordinated in time and space with cell cycle progression [33]. While ParABS was demonstrated to be essential for survival in a few bacterial species [34–37], its inactivation or depletion usually leads to pleiotropic phenotypes emanating from impaired *oriC* segregation, including aberrant chromosome numbers but also cell cycle progression and cell division defects [3].

ParABS systems translocate chromosomes in a variety of organisms with vastly different lifestyles, cell cycles, and differentiation programs. Despite significant advances in understanding the subcellular dynamics, biochemistry, and interactions of the ParABS components during chromosome segregation [3,38], the adaptation of this highly conserved system across species and between developmental stages remains unclear. Only a few investigations have examined the regulatory mechanisms tuning the ParABS system [39]. For example, studies have highlighted the importance of balancing the ParA-ParB levels to ensure correct cell cycle progression in *Caulobacter crescentus* [40], the ParA (Soj)-dependent transcriptional modulation of the *parAB* operon in *Bacillus subtilis* [41] and the developmental regulation of *parA* and *parB* expression that couples the segregation of the linear chromosome with the mycelial lifestyle of *Streptomyces coelicolor* [42]. Except for the latter, most available data on ParABS-dependent chromosome partitioning come from bacteria that proliferate classically through vegetative growth and binary division (one mother cell generating two daughter cells). However, many species possessing a ParABS system rely on relatively complex cell cycles involving non-binary division events, generating larger and sometimes variable numbers of progeny from a polyploid mother cell (*i.e., Cyanobacteria, Bdellovibrionata, and Actinobacteria*) [43–45]. The exploration of how the ParABS system partitions numerous copies of the chromosome during the intricate cell cycle of these organisms is still limited.

The obligate predatory bacterium *Bdellovibrio bacteriovorus* proliferates through filamentation and non-binary division inside the envelope of other diderm bacteria. Its cell cycle comprises two main phases determined by its presence outside or inside a prey [46,47]. The first phase, called attack phase (AP), corresponds to a G1 cell cycle stage during which *B. bacteriovorus* do not replicate their single circular chromosome as they search for prey [48]. Upon contact, predators attach to the prey surface and invade their periplasm, i.e., the space located between the inner and outer membranes [49,50]. Upon a G1-S transition, *B. bacteriovorus* starts the second stage of its cell cycle called the growth phase (GP), during which the cell elongates and copies its DNA multiple times. A first round of chromosome replication and segregation, which starts from the *oriC* localized at one cell pole [48], is quickly followed by the asynchronous firing of additional replication rounds using the *oriC* of multiple chromosome copies as template [48,51]. The newly synthesized chromosomal origins segregate progressively [48], unlike in *Streptomyces*, which also grows as polyploid filaments but uncouples DNA replication during vegetative growth from the ParABS-dependent segregation of multiple chromosome copies at sporulation. The filamentous and polyploid mother *B. bacteriovorus* cell eventually divides by synchronous constriction at multiple locations along the cell body, releasing a variable, odd or even number of offspring [48,51–53]. The extent of the GP and the number of predator daughter cells is determined by the size of the prey [54], which remains a closed nest as it is being digested by *B. bacteriovorus*, until newborn predators break the prey open to resume their predatory cycle [55].

In contrast to all other ParB homologs involved in chromosome segregation that always localize at the *parS* site [35,37,56–67], ParB in *B. bacteriovorus* does not mark the centromere throughout cell cycle. Our previous report showed that fluorescently tagged ParB, produced at native or constitutive levels, nucleates near *oriC* only after the initiation of chromosome replication, while the protein displays a diffuse localization in the predator progeny [48]. The constitutive production of ParB disturbs the progressive segregation of chromosomal copies, resulting in newborn *B. bacteriovorus* with aberrant cell length, *oriC* numbers, and nucleoid size. Thus, although ParB is critical for proper *oriC* partitioning and cell cycle progression in *B. bacteriovorus*, the formation of a ParB·*parS* higher order structure at the centromere is prevented during the non-proliferative stage of the predatory cycle [48]. The underlying mechanism behind this cell cycle-dependent on-off behaviour of ParB is unknown. Furthermore, the subcellular dynamics of ParA have not been explored in bacteria that segregate multiple pairs of sister chromosomes during their replicative phase of growth. An in-depth view of the ParABS system throughout the lifecycle of *B. bacteriovorus* is crucial to better understand how it operates in species with non-binary proliferation. With its intricate chromosome dynamics and unique centromere organization during the cell cycle, *B. bacteriovorus* represents a compelling model to uncover new levels of regulation in the highly conserved ParABS system.

Here we provide key insights into the ParABS system in *B. bacteriovorus*, by monitoring the subcellular localization, expression, and protein levels of ParA_Bb_ and ParB_Bb_ during all stages of the predatory cell cycle. Moreover, we characterize ParB_Bb_ CTPase and *parS* binding activities. Our data reveal multiple regulation layers in the ParABS system of this bacterium. We show that ParA and ParB protein levels fluctuate differently during the cell cycle despite being expressed from the same operon, their fine balance being critical for proper cell cycle progression. Uniquely, the genomic context of the *parS* sites during the non-replicative stage, combined with ParB_Bb_ protein depletion when the mother cell divides, play a critical role in the unique cell cycle-dependent organization of the centromere.

## Results

### The ParABS system is essential for B. bacteriovorus and exhibits specific localization patterns

The *B. bacteriovorus* genome encodes homologs of *parA* and *parB* and harbors two *parS* sites near the chromosomal origin, indicating the presence of a ParABS system for chromosome segregation **(Fig 1A)**. Our attempts to individually delete *parA_Bb_* (n = 86) and *parB_Bb_* (n = 90) were unsuccessful, underscoring the essential role of ParABS in this organism. We have shown previously that ParB_Bb_ labeled with the fluorescent protein mCherry localizes as foci that mark the multiple copies of chromosomal *oriC* during their partitioning in GP [48], consistent with the conserved role of ParB in chromosome segregation. However, the localization of ParB_Bb_ appeared to be cell-cycle regulated in *B. bacteriovorus* [48], whereas all ParB proteins described in other species permanently cluster at the centromere [35,37,56–67]. Using a ParB_Bb_-msfGFP fusion produced as a single copy from the native chromosomal locus, we confirmed that ParB_Bb_ forms foci only during the growth phase of the predator cell cycle **(Fig 1B, Video S1)**. Conversely, the ParB_Bb_-msfGFP fluorescence signal was weak and diffuse in the cytoplasm in non-replicating cells, i.e., in the free attack-phase cells (**Fig 1C**) or during the time window corresponding to the G1-S transition upon prey invasion [48] (**Fig 1B)**.

**Fig 1.**
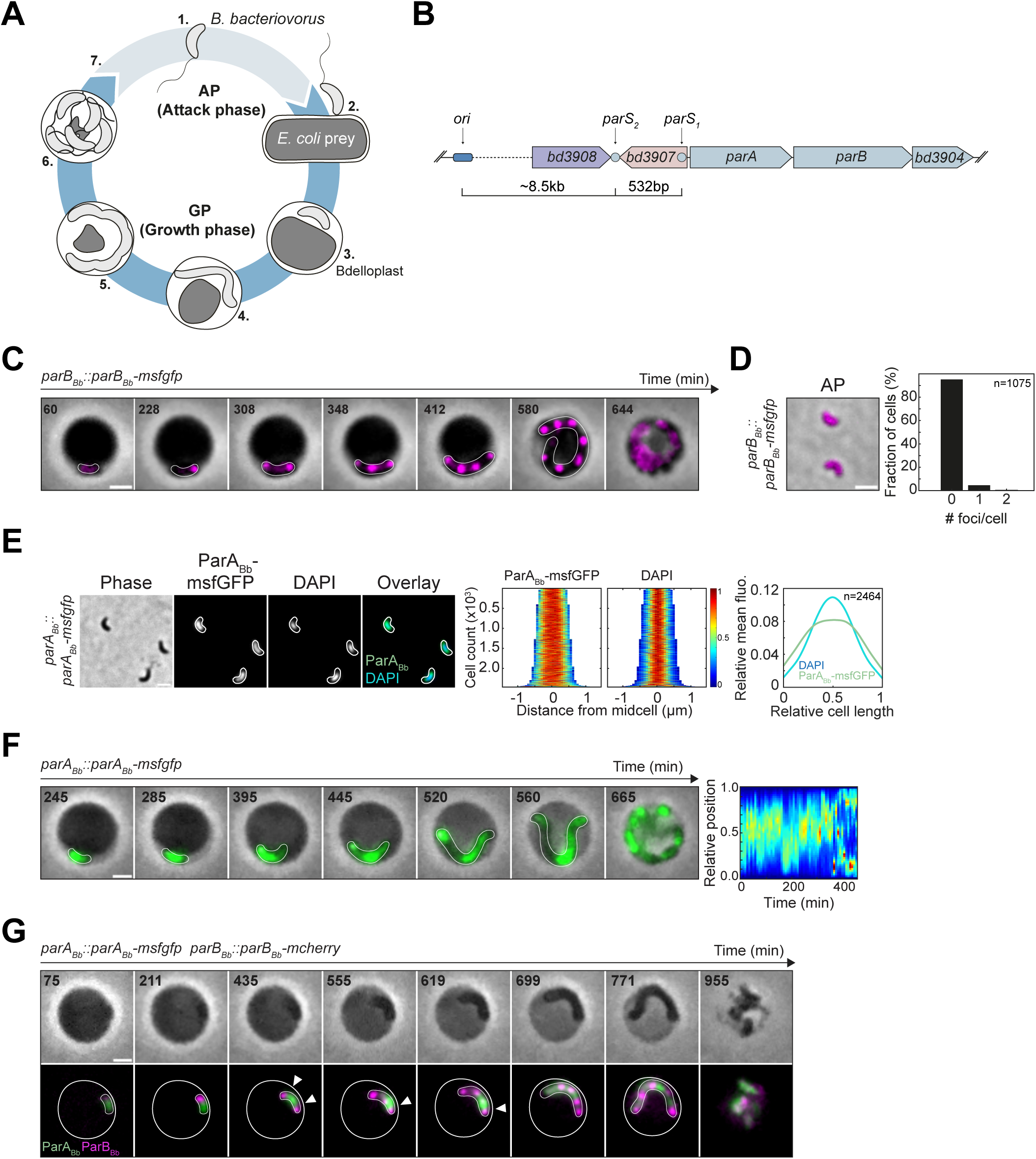
Subcellular localization of the ParABS system during the *B. bacteriovorus* cell cycle. (A) Schematics of the *Bdellovibrio bacteriovorus* cell cycle. Numbers indicate key steps in the cycle: 1. Freely-swimming attack phase (AP) cells, 2. Attachment of *B. bacteriovorus* to its prey, 3. *B. bacteriovorus* resides in the periplasm of the prey, which is now called bdelloplast, 4. Filamentous growth and consumption of prey content, 5. Pre-divisional state, 6. The non-binary division of the mother cell generates an odd or even number of daughter cells, which mature before 7. escaping the prey remnants and resuming the cell cycle. Attack phase (AP) and growth phase (GP) represent the non-replicative and replicative stages of the cell cycle, respectively. (B) Chromosomal context of the *parABS* system in the WT *B. bacteriovorus* genome. The *parAB* operon is shown in blue. *bd3908* encodes a putative rRNA methyltransferase*; bd3907* encodes an unknown protein*; parA* and *parB* are *bd3906* and *bd3905,* respectively; *bd3904* encodes a bactofilin homolog. The genomic distance between chromosomal *ori* and both *parS* sites is indicated. (C) Cell-cycle dependent localization of ParB_Bb_. *B. bacteriovorus* strain *parB_Bb_::parB_Bb_-msfgfp* (GL1654) was mixed with *E. coli* prey and imagined in time-lapse after 60 min with 8-min intervals. Overlays of phase contrast and fluorescence images of selected time points are shown; multiple well-separated ParB_Bb_-msfGFP foci appear over time; in this example, ParB_Bb_-msfGFP foci disappear at time point 664 min. The time-lapse is shown in Video S1. (D) Endogenous ParB_Bb_ does not form an *oriC?*-bound focus during AP. Overlay of phase contrast and fluorescence images of the *parB_Bb_::parB_Bb_-msfgfp* strain (GL1654) for representative AP cells; histogram representing the percentage of cells with zero, one, or two ParB_Bb_-msfGFP foci in the same strain. (E) ParA_Bb_ is nucleoid bound. Left to right: representative phase contrast and fluorescence images of AP cells of the *parA_Bb_::parA_Bb_-msfgfp* strain (GL2134) stained with DAPI; demographs of the corresponding fluorescent signals in the same cells sorted by length and oriented based on signal intensity; heatmaps represent relative fluorescence intensities; mean pole-to-pole profiles of relative fluorescence intensity of the corresponding signals in the same cells. (F) Endogenous ParA_Bb_ moves dynamically along the growing predator filament. From left to right: *B. bacteriovorus* strain *parA_Bb_::parA_Bb_-msfgfp* (GL2134) was mixed with prey and imaged in time-lapse after 80 min with 5-min intervals; left: overlay of phase contrast and fluorescence images, time points are shown in min; right: kymograph of the ParA_Bb_-msfGFP signal along the cell length for the same cell until time point 445 min. The time-lapse is shown in Video S2. (G) The dynamic interplay between ParB_Bb_ and ParA_Bb_. *B. bacteriovorus* strain *parA_Bb_::parA_Bb_-msfgfp parB_Bb_::parB_Bb_-mcherry* (GL2154) was mixed with prey and imagined in time-lapse after 75 min with 8-min intervals. Top: phase contrast; bottom: overlay of ParB_Bb_-mCherry and ParA_Bb_-msfGFP signals. Arrowheads point to a ParB_Bb_ focus on the edge of a ParA_Bb_-msfGFP cloud. The time-lapse is shown in Video S3. Scale bars are 1 µm. n indicate the number of cells analyzed in a representative experiment. *B. bacteriovorus* and bdelloplasts outlines in panels C, F, G were drawn manually based on the phase contrast images. All experiments were performed at least twice. See also Fig S1.

To obtain more insights into the functioning of the ParABS system during the cell cycle of *B. bacteriovorus*, we constructed a functional ParA_Bb_-msfGFP fusion **(Fig S1A)** produced natively as a single copy and monitored its subcellular localization. We found that ParA_Bb_-msfGFP co-localized with the nucleoid in non-replicative cells (**Fig 1D**), in agreement with the non-specific DNA binding of ParA proteins [16,17]. Unlike in other species with a polarly localized *oriC*, ParA_Bb_-msfGFP localization was not visibly biased towards a particular cell pole, which we were able to distinguish using the invasive pole marker RomR-mCherry [48,68] **(Fig S1B**). During the *B. bacteriovorus* proliferative phase inside the prey, ParA_Bb_-msfGFP displayed a dynamic localization pattern reminiscent of the DNA-bound ParA “cloud” characterized in other organisms (**Fig 1E, Video S2**), which pulls apart duplicated ParB·*parS* partitioning complexes via repeated ParA-ParB interactions [19,69]. In addition, we detected an interaction between ParA_Bb_ and ParB_Bb_ in a bacterial two-hybrid assay (**Fig S1C**), suggesting a similar interplay between these proteins in *B. bacteriovorus*. However, in the growing filament, several ParA_Bb_-msfGFP accumulations were present at the same time and moved dynamically along the filamentous predator cell, shifting from one subcellular region to another (**Fig 1E, S2 Movie**). Using a strain carrying both fluorescently labeled ParA_Bb_ and ParB_Bb_ (**Fig S1D**), we observed that ParB_Bb_-mCherry foci were mainly located at the edges of ParA_Bb_-msfGFP clouds **(Fig 1F, arrowheads, Video S3)**. This complex localization pattern is consistent with the idea that the *B. bacteriovorus* ParABS system is adapted to drive the simultaneous or sequential segregation of multiple chromosome copies.

### A fine-tuned balance of parA_Bb_ and parB_Bb_ expression underlies proper chromosome segregation and cell cycle progression in B. bacteriovorus

Remarkably, the ParB_Bb_-associated fluorescence seemed to drop at the end of the cell cycle, unlike ParA_Bb_-msfGFP (see the last time points in **Fig 1B, 1E**). Thus, our data suggests that these key players of the ParABS system undergo an unknown regulation resulting in distinct changes in protein abundance throughout the predatory cell cycle. To delve deeper into the cell-cycle control of the ParABS system in *B. bacteriovorus*, we first investigated its transcriptional regulation. RT-PCR **(Fig 2A)** shows that *parA_Bb_* and *parB_Bb_* are part of the same operon. Furthermore, their expression is cell-cycle regulated as the corresponding RNA was detected mainly in late GP and not in AP **(Fig 2A)**, in agreement with previous data including RNAseq [70,71]. Consistent with the significance of the cell cycle-dependent *parAB_Bb_* expression in *B. bacteriovorus,* we previously found that the constitutive expression of *parB_Bb_* (*parB*_*Bb*_^++^) negatively impacts cell cycle progression and chromosome segregation [48]. Altering the native *parA_Bb_* expression similarly impaired cell cycle progression and *ori* partitioning, mimicking the *parB*_*Bb*_^++^ phenotype (**Fig 2B, Fig S2A-B**). Remarkably, the constitutive, simultaneous expression of both *parA_Bb_* and *parB_Bb_* (from the same promoter) decreased the severity of the single overexpression phenotypes (**Fig 2B, Fig S2A-C**). Thus, our data show that the ParABS system is under transcriptional control during the *B. bacteriovorus* cell cycle and that this level of regulation is crucial for proper chromosome segregation and cell cycle progression, likely by contributing to a fine balance between both partners.

**Fig 2.**
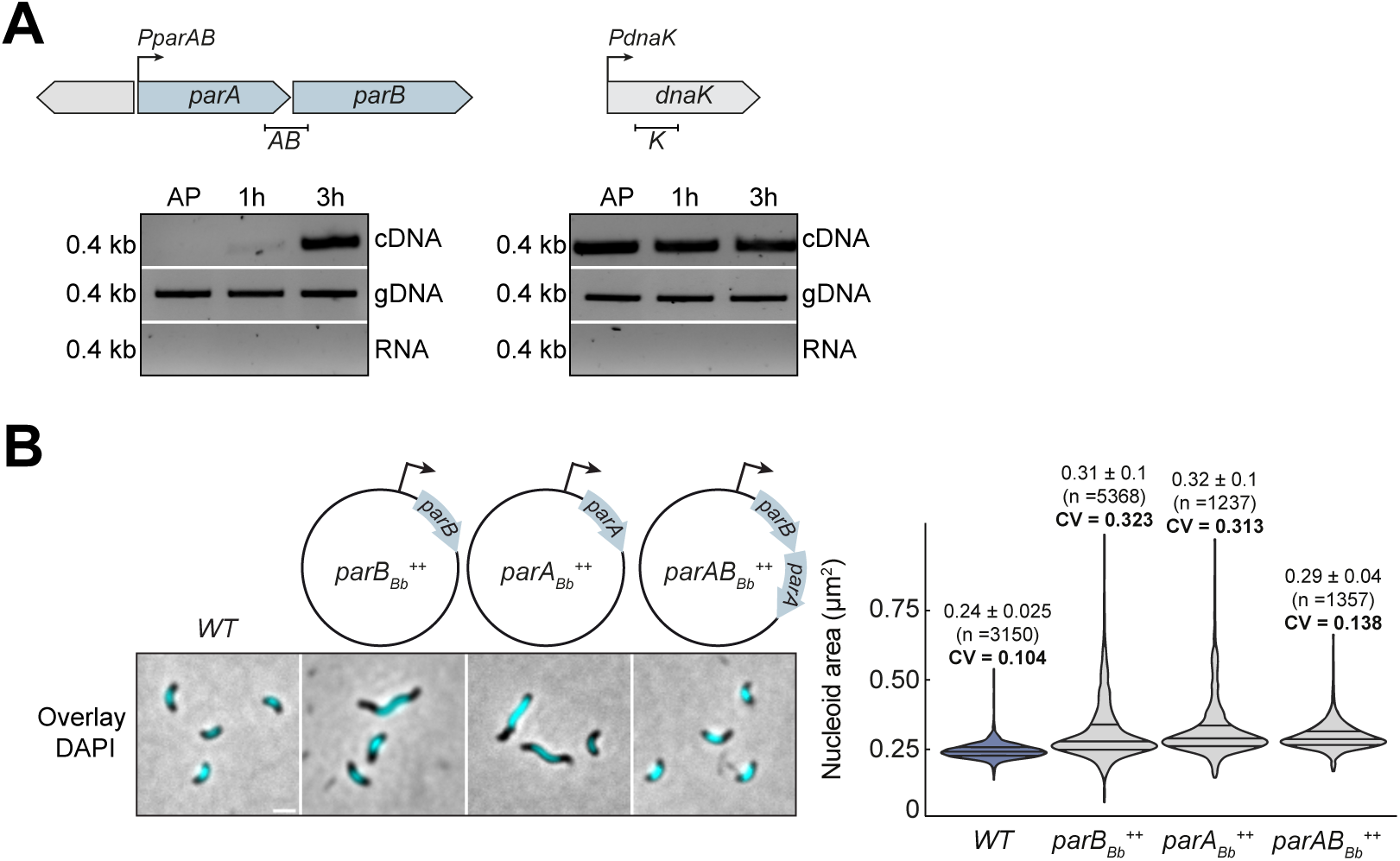
Biphasic expression of the *parAB* operon during the *B. bacteriovorus* cell cycle is critical for proper chromosome segregation. (A) Biphasic expression of the *parAB_Bb_* operon. Top: schematic representation of the *B. bacteriovorus parAB* and *dnaK* (housekeeping gene) genomic context. The fragments corresponding to the *parA-parB* junction or inside the *dnaK* gene amplified by RT-PCR are indicated below (*AB,* 400 bp; *K*, 400 bp). RNA samples were isolated from wild-type *B. bacteriovorus* in AP, 1 h, and 3 h after mixing with prey. Bottom: agarose gel analysis of the indicated fragments amplified by PCR from cDNA (top panel), genomic DNA (middle panel, positive control), and total RNA (bottom panel, negative control). Molecular weight markers (in kb) are indicated on the left. (B) Balanced *parA_Bb_:parB_Bb_* expression is essential for proper chromosome partitioning. Left: phase contrast and fluorescence images of representative AP cells stained with DAPI. The schematics illustrate the presence of a plasmid allowing the constitutive expression (from the P*nptII* promoter) of the indicated gene(s) *parB_Bb_* (GL1261), *parA_Bb_* (GL1460), or both (GL1004) in an otherwise *WT* background. Cells with abnormal nucleoid areas are observed when *parA_Bb_* (*parA_Bb_^++^*) or *parB_Bb_* (*parB_Bb_ ^++^*) is overexpressed, but to a lower extent when both are overexpressed (*parAB_Bb_ ^++^*). Scale bar, 1µm. Right: violin plot of cell length, cell area, and nucleoid area distributions in the same cells. The lines indicate the 25, 50, and 75 percent quantiles from bottom to top. Mean, standard deviation and coefficient of variation (CV) values are shown on top of the corresponding plot. n indicate the number of cells analyzed in a representative experiment. All experiments were performed at least twice. See also Fig S2.

### The protein levels of ParA_Bb_ and ParB_Bb_ vary differently during the predatory cell cycle

Since ParA_Bb_ and ParB_Bb_ fluorescence profiles hinted at distinct relative protein abundance despite identical transcriptional regulation, we sought to obtain detailed insights into ParA_Bb_ and ParB_Bb_ protein levels during the cell cycle. Western blot using an antibody against ParB_Bb_ shows that the endogenous (untagged) ParB_Bb_ protein levels vary during the cell cycle, being almost unnoticeable in AP but present in higher amounts in GP (**Fig 3A, S3A**), mirroring the RNA levels (**Fig 2A**) and the fluorescence signal of the natively produced ParB_Bb_-msfGFP (**Fig 1B**). Accordingly, single-cell analysis of the ParB_Bb_-msfGFP fluorescence intensity profile over time indicates that ParB_Bb_ proteins accumulate during cell growth but rapidly drop when *B. bacteriovorus* divides into multiple daughter cells (**Fig 3B, magenta, Fig S3B**). Notably, the levels of natively produced ParB_Bb_-msfGFP or ParB_Bb_-mCherry protein fusions, detected with the same anti-ParB_Bb_ antibody, fully reflected the endogenous untagged ParB_Bb_ profile during a synchronized *B. bacteriovorus* cell cycle **(Fig S3C)**, confirming that these fusions are reliable reporters of the native ParB_Bb_ protein.

**Fig 3.**
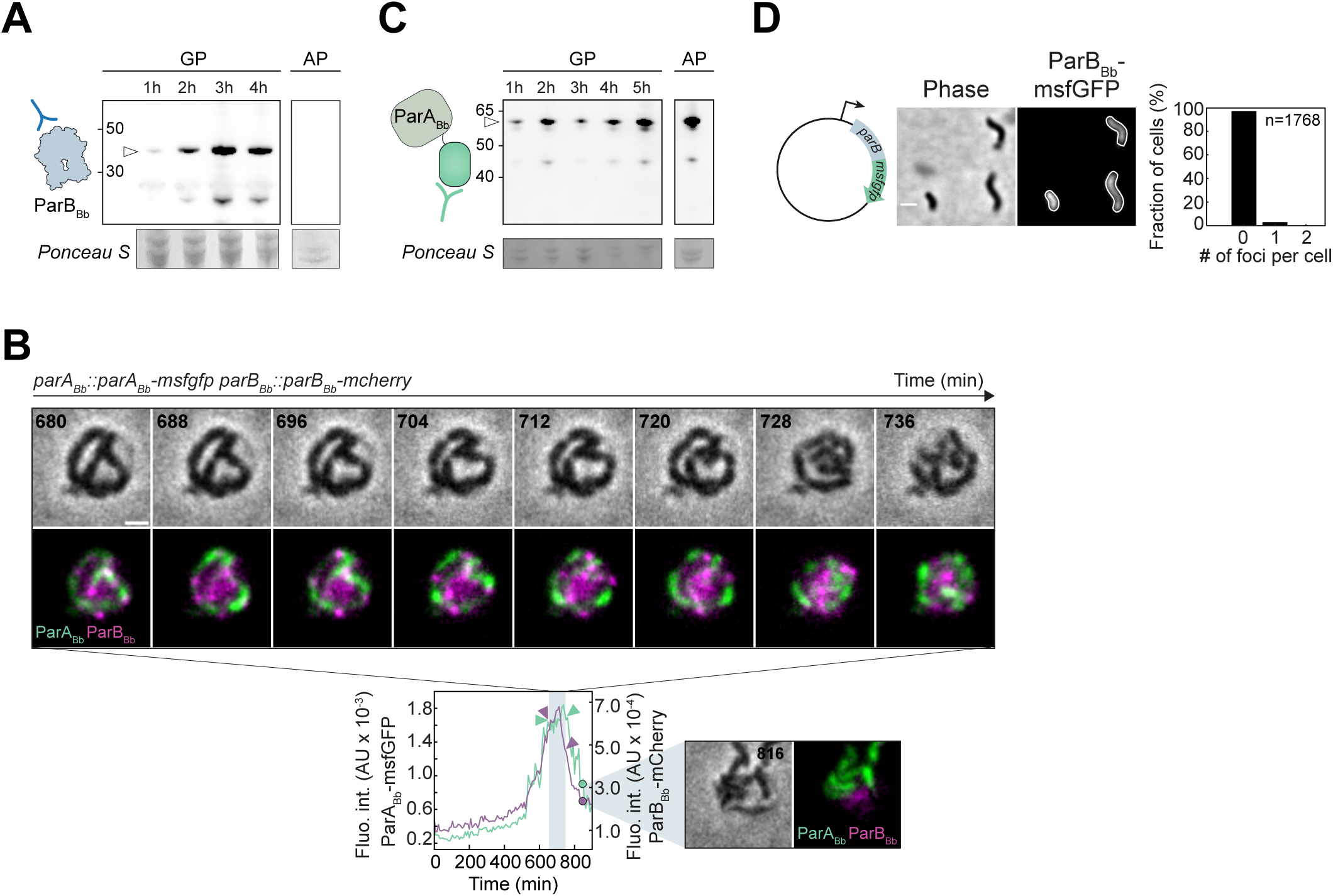
The levels of ParB_Bb_ and ParA_Bb_ vary differently during the cell cycle. (A) ParB_Bb_ protein levels are cell-cycle dependent. Western blots of whole-cell protein extracts from *WT B. bacteriovorus* were probed with an anti-ParB_Bb_ antibody, as represented in the schematics on the left. WT protein samples are isolated at time points throughout the predatory cell cycle: AP, 1h, 2h, 3h, and 4h after mixing with prey (GP). Arrowhead points to ParB_Bb_ (detected only during GP). Ponceau staining of the same membranes (where bands were most visible, ∼30-50 kDa) is shown below each blot as a loading control. Molecular weight markers (kDa) are shown on the side. (B) ParB_Bb_-msfGFP fluorescence intensity level drops during cell division while ParA_Bb_-msfGFP fluorescence intensity levels remain high. *B. bacteriovorus* strain *parA_Bb_::parA_Bb_-msfgfp parB_Bb_::parB_Bb_-msfgfp* (GL2154) was mixed with prey and imaged in time-lapse after 75 min with 8-min intervals. Top: overlay of phase contrast and fluorescence images for ParA_Bb_-msfGFP and ParB_Bb_-mCherry for selected late time points; Bottom: mean fluorescence intensities of msfGFP (in green) and mCherry (in magenta) plotted over time for the same cell. The time window corresponding to the images displayed on top is represented as a vertical blue rectangle. ParB_Bb_-mCherry fluorescence intensity decreases during this interval, while ParA_Bb_-msfGFP fluorescence decreases later when daughter cells start leaving the prey. A representative later time point is shown on the right. (C) ParA_Bb_ protein levels do not decrease in the attack phase. Western blots of whole-cell protein extracts from *parA_Bb_::parA_Bb_-msfgfp B. bacteriovorus* (GL2134) were probed with an anti-msfGFP antibody, as represented in the schematics on the left. GL2134 protein samples are isolated at time points throughout the predatory cell cycle: 2h, 4h, 6h after mixing with prey and AP. Arrowhead points to ParA_Bb_-msfGFP. Free msfGFP constitutively produced in *B. bacteriovorus* is shown as control (GL1212). Ponceau staining of the same membranes (where bands were most visible, ∼30-50 kDa) is illustrated below each blot as a loading control. Molecular weight markers (kDa) are shown on the side. (D) Overproduction of ParB_Bb_-msfGFP does not lead to the *parS-*bound focus in AP. Left: representative phase contrast and fluorescence images of AP cells of a WT *B. bacteriovorus* strain constitutively producing ParB_Bb_-msfGFP (GL1003). Right: histogram representing the percentage of cells with zero, one, or two ParB_Bb_-msfGFP foci in the same strain; n indicates the number of cells analyzed in a representative experiment. Scale bars are 1 µm. All experiments were performed at least twice. See also Fig S3.

Unlike ParB_Bb_, ParA_Bb_-msfGFP was detected throughout the cell cycle, including at the end of GP and in AP, where protein levels were higher (**Fig 3C**), consistent with the corresponding fluorescence signal (**Fig 1E**). ParA_Bb_-msfGFP fluorescence intensity remained high during the last cell cycle stages, including cell division (**Fig 3B, green, Fig S3D**). These results show that, besides cell-cycle regulated transcription, the ParABS system in *B. bacteriovorus* is subjected to an additional level of control that results in distinct ParA_Bb_ and ParB_Bb_ protein levels at different cell cycle stages. While *in vitro* studies showed that the ParA:ParB balance is important for efficient partitioning [72,73], this is the first evidence of cell-cycle-dependent changes in the relative levels of ParA and ParB proteins.

### The formation of ParB**_Bb_**·parS complexes is inhibited in non-replicative predator cells

In addition to the sudden decrease of ParB_Bb_ protein levels at the end of the GP, the subcellular localization of ParB_Bb_ fluorescent fusions also changed, shifting from clear foci to diffuse localization **(Fig 1B, 3B)**. However, both events could not be easily distinguished due to their occurrence within a relatively short time window and the low fluorescence of the natively produced ParB_Bb_ fusion at that stage. Therefore, we sought to uncouple the ability of ParB_Bb_ to form foci (due to *parS* binding and spreading on adjacent DNA) from ParB_Bb_ protein levels by constitutively expressing *parB_Bb_-msfgfp*. In these cells, the fluorescence signal **(Fig S3E)** and protein levels (**Fig S3F)** of ParB_Bb_-msfGFP in AP were stronger than upon expression from the native locus. Still, overproduced ParB_Bb_-msfGFP was diffuse in AP cells **(Fig 3D)**, consistent with our previous results with ParB_Bb_-mCherry [48]. Furthermore, these cells displayed the same phenotypes observed when the untagged ParB_Bb_ is constitutively produced (i.e., morphological and chromosome segregation defects; **Fig S3G, Fig S2B)**, further supporting the relevance of these ParB_Bb_ fusions. Thus, a third level of control is applied to ParB in *B. bacteriovorus* to prevent its accumulation at the centromere, regardless of its protein level.

### ParB_Bb_ specifically accumulates on B. bacteriovorus parS in a CTP-dependent manner

The formation of partitioning complexes in other species requires both the initial nucleation and the spreading of ParB on the DNA [8,10,11,13,14,74]. To investigate ParB_Bb_’s ability to perform these two critical functions, we assessed its capacity to bind and spread from *parS* and to hydrolyze CTP *in vitro.* The two *parS* sequences in the proximity of the *B. bacteriovorus* chromosomal *ori* are identical and closely resemble the *parS* consensus **(Fig S4A)** [2]. In a biolayer interferometry assay, purified ParB_Bb_ bound a linear DNA substrate carrying *parS_Bb_*, and the addition of CTP triggered ParB_Bb_ sliding off (marked by the higher Kd in presence of CTP; **Fig 4A**), like other characterized ParB homologs [11,13– 15,74]. Moreover, ParB_Bb_ accumulated on a closed DNA loop containing *parS_Bb_* in a CTP-dependent manner, reflecting the capacity of ParB_Bb_ to nucleate on *parS_Bb_* and escape onto neighboring DNA **(Fig 4B)**. This accumulation was specific to the cognate *parS_Bb_*, as no binding was observed when we used *parS* from *Caulobacter crescentus* (*parS_Cc_;* **Fig 4B)**. Finally, measurements of inorganic phosphate release showed that ParB_Bb_ hydrolyses CTP but no other NTPs in the presence of *parS_Bb_* **(Fig S4B)**. Thus, the *B. bacteriovorus* ParB protein is a bona-fide *parS*-binding CTPase. Consistent with those *in vitro* data, ParB_Bb_ specifically accumulated on *parS_Bb_ in vivo*, using *E. coli* (which lacks an endogenous ParABS) as a heterologous system. Indeed, ParB_Bb_-msfGFP formed foci when produced in *E. coli,* only in the presence of a plasmid carrying *parS_Bb_* and not when the plasmid carried the non-cognate *parS_Cc_* (**Fig 4C, Fig S4C**). Interestingly, ParB from *Caulobacter crescentus* also failed to form a focus and was diffuse in the cytoplasm when ectopically produced in AP *B. bacteriovorus* cells **(Fig 4D)**, even though ParB_Cc_ can bind *parS_Bb_ in vitro* **(Fig S4D)** and in *E. coli* **(Fig S4E)**. Therefore, the absence of ParB·*parS* complex in non-replicative *B. bacteriovorus* cells is unlikely to result from an unusual functionality of the *B. bacteriovorus* ParB protein itself.

**Fig 4.**
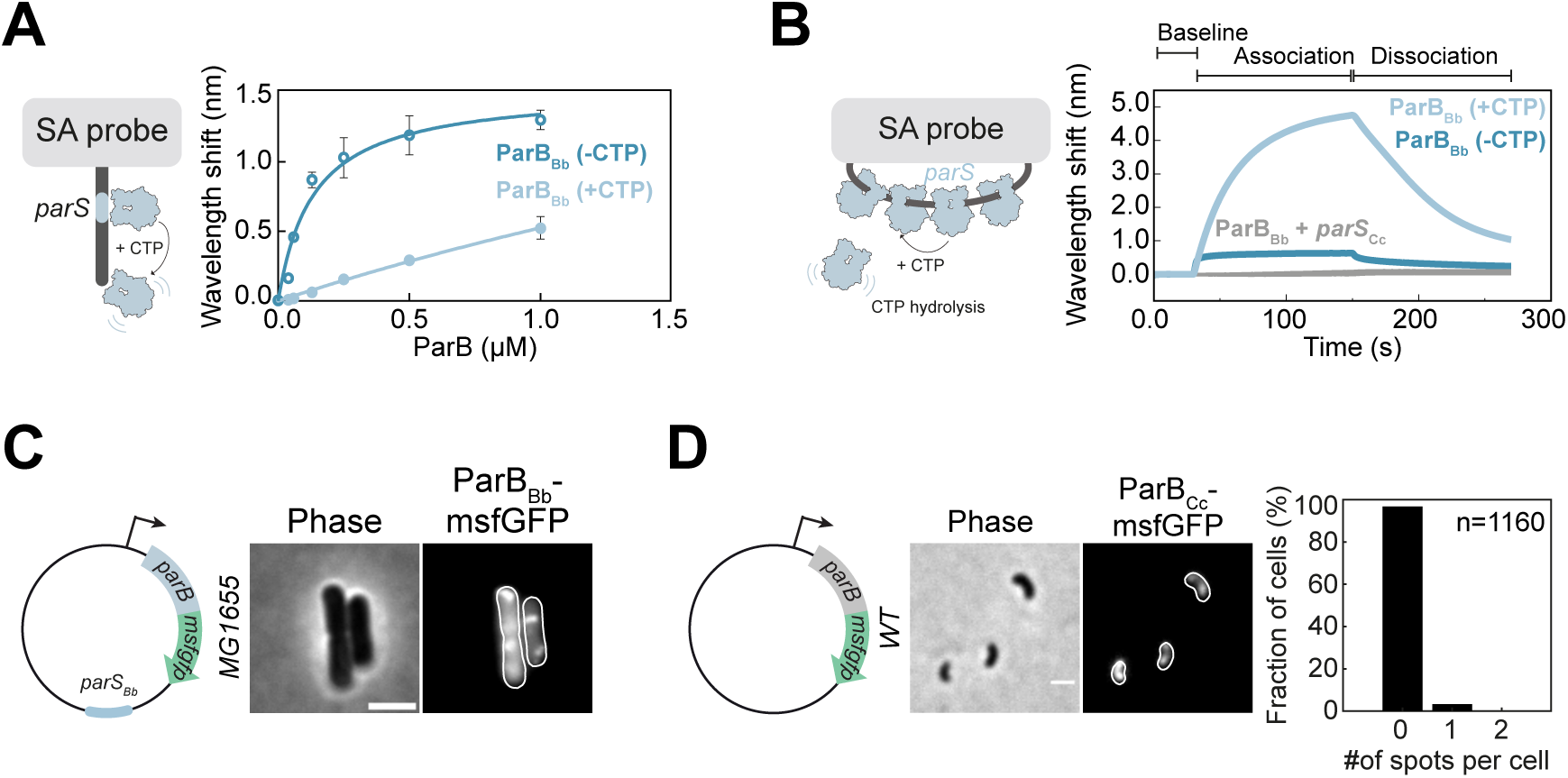
ParB_Bb_ is a CTPase with a specific affinity for its cognate *parS_Bb_*. (A) CTP reduces the nucleation of ParB_Bb_ on *parS_Bb_.* Left: schematics of the Streptavidin (SA)-coated probe and the linear 40-bp DNA substrate containing the cognate *parS_Bb_* sequence. Right: ParB_Bb_ titration curves in the presence or absence of CTP obtained from biolayer interferometry (BLI) analysis (see B and Material and Methods). All reactions contained 1 mM CTP (when appropriate), 0.5 µM 40 bp *parS_Bb_* DNA, and an increasing concentration of ParB_Bb_. The BLI experiment was done in triplicates; data were fitted to calculate the binding affinity constant Kd (nM); Kd = 134 ± 23 nM for the 40 bp *parS_Bb_*, and Kd = 5500 ± 397 nM for the 40 bp *parS_Bb_* + 1mM CTP. (B) ParB_Bb_ accumulation at *parS_Bb_* is specific and CTP-dependent. Left: schematics of the SA-coated probe and a closed DNA substrate with a *parS* sequence, allowing ParB accumulation through nucleation and CTP binding-dependent spreading. CTP hydrolysis removes ParB from the DNA. Right: BLI analysis of the interaction between 1 µM ParB_Bb_-6xhis in the presence or absence of 1 mM CTP and a 180 bp DNA substrate containing the cognate *parS_Bb_* (blue) or non-cognate *parS_Cc_* (grey) sequence. The BLI probe was dipped into a buffer-only solution (base), then into a premix of protein +/-CTP (association), and finally returned to a buffer-only solution (dissociation). Each BLI experiment was done in triplicate, and a representative sensorgram is presented. (C) ParB_Bb_ protein can bind to its cognate *parS* sequence in *E. coli*. Representative phase contrast and fluorescence images of *E. coli* carrying a plasmid with *parS_Bb_* and allowing constitutive production of ParB_Bb_-msfGFP (GL1669; plasmid schematics on the left). The scale bar is 2 µm. (D) A constitutively produced fusion of *Caulobacter crescentus* ParB (ParB_Cc_) to msfGFP does not form foci in AP *B. bacteriovorus* cells. Left: representative phase contrast and fluorescence images of AP cells of a WT strain constitutively producing ParB_Cc_-msfGFP from a plasmid (GL2108; schematics on the left). The fluorescence signal shows the partial nucleoid exclusion pattern of freely diffusing cytosolic proteins, characteristic of AP *B. bacteriovorus* cells ^1^. Right: histogram of the percentage of cells with zero, one, or two ParB_Cc_-msfGFP foci in the same strain; n indicates the number of cells analyzed in a representative experiment. The scale bar is 1 µm. See also Fig S4.

### The parS_Bb_ chromosomal context prevents ParB_Bb_ accumulation in non-replicative cells

Having established that the ParB_Bb_ activities are not *per se* drastically different than previously characterized ParB homologs, we investigated the possibility that in AP cells, the chromosomal context of the centromere is incompatible with ParB_Bb_ accumulation. Moving one or both *parS_Bb_* sites to a remote location on the chromosome is expected to result in pleiotropic effects due to a major perturbation of segregation, which would prevent unambiguous interpretation. Instead, we addressed whether the genomic context impacts ParB_Bb_ localization by placing a *parS_Bb_* site on a replicative plasmid and constitutively producing ParB_Bb_-msfGFP from the same vector in *B. bacteriovorus* **(Fig 5A)**. Remarkably, we found that ParB_Bb_-msfGFP localizes as clear foci in AP cells only when the plasmid carries the intact *parS_Bb_.* No focus was observed when we used a mutated *parS_Bb_\** as a control **(Fig S5A)**. These results indicate that in AP cells, ParB_Bb_ has the capacity to bind *parS_Bb_* specifically but only when *parS_Bb_* is out of its native chromosomal context. Moreover, AP *B. bacteriovorus* cells carrying the *parS_Bb_* + *parB_Bb_-msfgfp^++^* vector did not exhibit the sick phenotype observed in *parB ^++^* cells **(Fig S5B**, compared with **Fig S2B)**, presumably because the plasmidic *parS_Bb_* titrates the excess ParB_Bb_ present in the cytoplasm, thereby alleviating the negative effect associated with its overproduction. Hence, our data show that the chromosomal context is an essential factor underlying the unique subcellular distribution of ParB_Bb_ in *B. bacteriovorus*.

**Fig 5.**
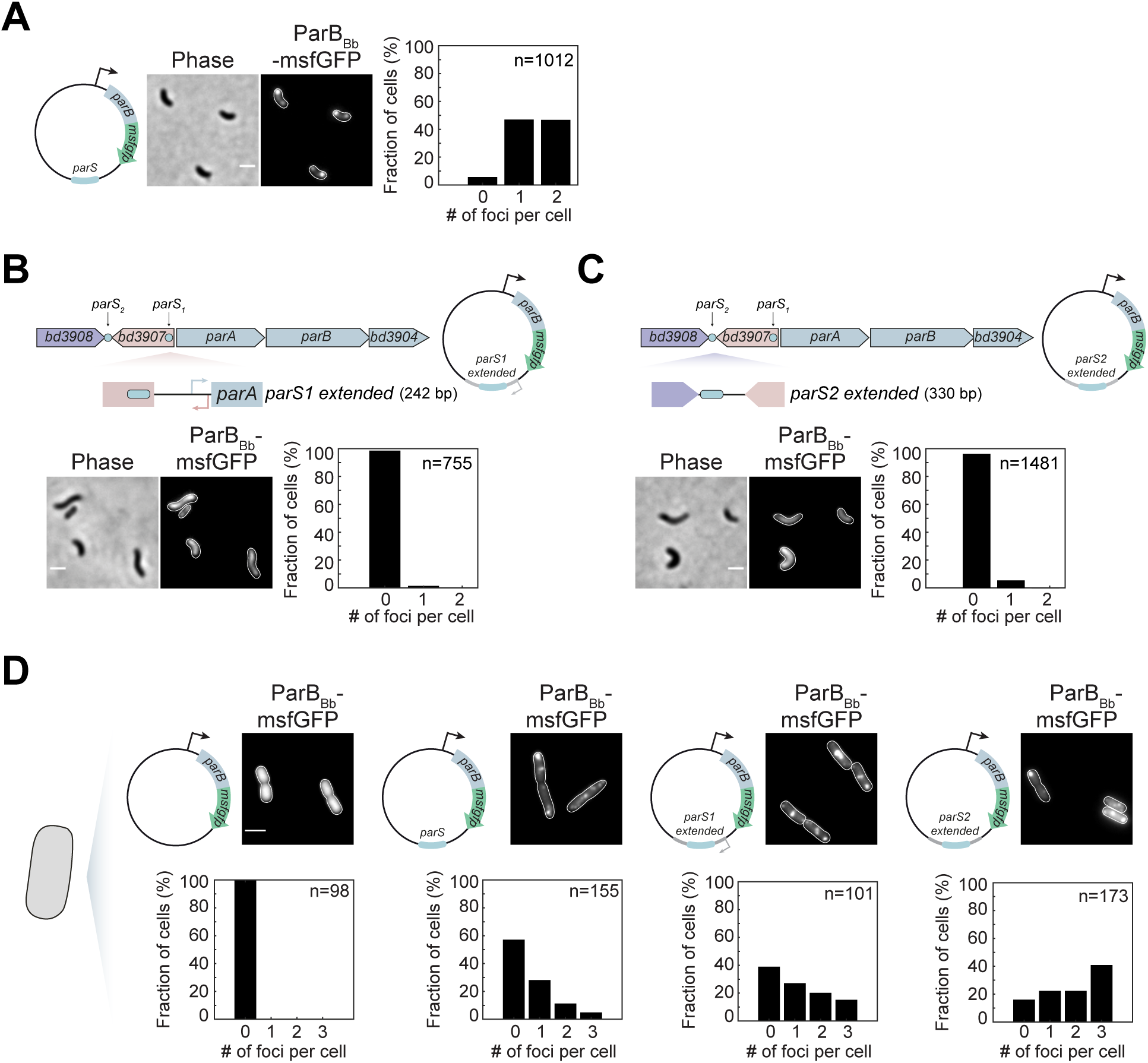
The genomic context of *parS_Bb_* plays a role in ParB_Bb_ nucleation. (A) Changing the native context of the *parS_Bb_* enables ParB_Bb_ binding. Left: representative phase contrast and fluorescence images of AP cells of WT *B. bacteriovorus* strain constitutively producing ParB_Bb_-msfGFP from a plasmid carrying a copy of the *parS_Bb_* sequence (GL1751). Right: histogram representing the percentage of cells with zero, one, or two ParB_Bb_-msfGFP foci in the same strain. (B-C) Short chromosomal regions around each *parS_Bb_* site are enough to prevent ParB_Bb_ from clustering on the centromere. Top: schematic representation of the *parAB* operon in *B. bacteriovorus* and the upstream *parS* sequences; chromosomal regions around *parS_Bb_* sequences (“extended *parS”*) cloned in the plasmids are indicated; plasmid schematics are shown on the right. Bottom: representative phase contrast and fluorescence images of AP cells of WT strain constitutively producing ParB_Bb_-msfGFP from a plasmid carrying *parS_1extended_* (B, GL1749) or *parS_2extended_* (C, GL1925); histogram representing the percentage of cells with zero, one, or two ParB_Bb_-msfGFP foci in the same strain. Scale bars are 1 µm. (D) The genomic context of the *parS_Bb_* prevents ParB_Bb_ nucleation only in *B. bacteriovorus.* Top: representative phase contrast and fluorescence images of *E. coli* strains carrying plasmids as in Figs 3D, 5A-C, constitutively producing ParB_Bb_-msfGFP in the absence of *parS* (GL1661), or in the presence of *parS_Bb_* (GL1669), *parS_1extended_* (GL1737) or *parS_2extended_* (GL1899), respectively. Schematics illustrate the *parB_Bb_–msfgfp* expression plasmid with or without (extended) *parS* sequences. Bottom: histogram representing the percentage of cells with zero, one, or two ParB_Bb_-msfGFP foci in the same strains. The scale bar is 1 µm. n indicates the number of cells analyzed in a representative experiment; all experiments were performed at least twice. See also Fig S5.

To obtain cues on the difference between the chromosomal and plasmidic *parS_Bb_* leading to distinct ParB_Bb_ localization patterns, we tested the impact of the *parS_Bb_* surrounding regions. We constructed two plasmids carrying the *parS_Bb_* sequence flanked by their short upstream and downstream chromosomal regions (named here *parS1 extended* and *parS2 extended*, spanning 242 bp and 330 bp, respectively, **Fig 5B-C**). Strikingly, the overproduced ParB_Bb_-msfGFP failed to form a fluorescent focus in both cases **(Fig 5B-C)**. Instead, the signal was diffuse, and cells exhibited the characteristic *parB_Bb_^++^* phenotype **(Fig S5B**, compared with **Fig S2B)**. This effect was limited to *B. bacteriovorus* since in *E. coli*, ParB_Bb_-msfGFP localized as foci despite the addition of *parS1* or *parS2 extended* regions **(Fig 5D)**. Therefore, our results suggest that a *Bdellovibrio*-specific factor makes the chromosomal environment of *parS_Bb_* sites incompatible with ParB_Bb_ clustering during the non-proliferative phase of the cell cycle.

## Discussion

Altogether, our study provides novel insights into the complex regulation of the conserved ParABS system in a bacterium that segregates multiple copies of its chromosome during its non-binary proliferation. We followed the subcellular dynamics, expression, and protein levels of the key players in that system, the proteins ParA and ParB, throughout the predatory cell cycle of *B. bacteriovorus*. Additionally, we assessed the ability of ParB_Bb_ to form nucleoprotein complexes on the centromere *in vitro* and *in vivo.* We discovered that the ParABS system is coupled to cell cycle progression via three levels of regulation that modulate (i) *parA_Bb_* and *parB_Bb_* gene expression, (ii) ParA_Bb_ and ParB_Bb_ protein levels, and (iii) chromosomal *parS* accessibility during the non-proliferative and the replicative stages of the predator lifecycle (**Fig 6**). Furthermore, our data indicate that the combination of these layers of cell cycle-dependent control contributes to clearing the *B. bacteriovorus* cell and the centromere from ParB_Bb_ during the non-proliferative stage, which is critical for proper cell proliferation.

**Fig 6.**
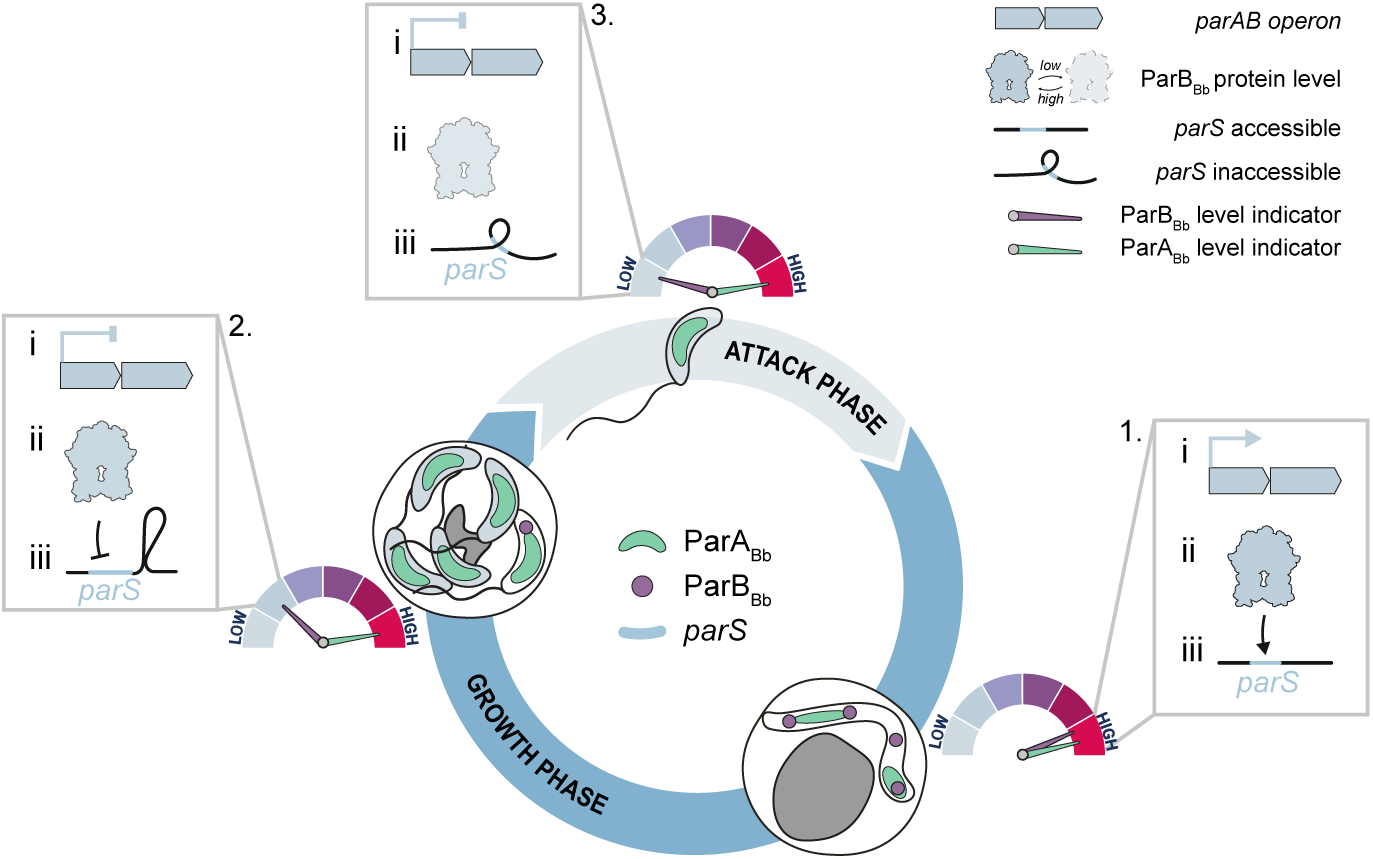
Model for coupling cell cycle progression and centromere organization through multi-layered regulation of the ParABS system in *B. bacteriovorus*. Schematic representation of the subcellular localization of ParA_Bb_ (green) and ParB_Bb_ (magenta) throughout the predatory cell cycle in *B. bacteriovorus*. The three levels of ParABS regulation are i) transcription of the *parAB* operon, ii) ParA_Bb_ and ParB_Bb_ protein levels, and iii) accessibility of the chromosomal *parS* site. In growing cells (1), the *parAB* operon transcription is “on”, ParB_Bb_ and ParA_Bb_ levels are high, and *parS* is accessible, enabling ParB_Bb_ to form nucleoprotein complexes on the centromere. The ParABS system rearranges concomitantly with non-binary division (2), transitioning towards an “off” state. During the attack phase (3), ParA_Bb_ levels remain high, but ParB_Bb_ levels are low due to potential protein degradation at the end of the growth phase, combined with the absence of new transcription of the *parAB* operon. The *parS* chromosomal context is incompatible with the formation of a ParB_Bb_ complex at the centromere during this stage of the cell cycle.

### Intricate subcellular dynamics of ParA and ParB during the replicative stage of the cell cycle

We investigated the localization of ParA in *B. bacteriovorus* during the predatory cell cycle using a functional, natively produced fluorescent fusion. The subcellular localization of ParA has been observed in *Streptomyces* species, which also feature a polyploid growth stage [75,76]. However, in these bacteria, the chromosome is copied multiple times during vegetative growth, while the ParABS system only acts later in a distinct developmental stage to separate nucleoids in future spores [77]. Thus, our data provide the first visualization of the ParA protein in a bacterium that segregates multiple copies of its chromosome during the replicative stage of its cell cycle. In non-replicative cells, ParA_Bb_ is uniformly distributed on the nucleoid, in contrast with the ParA gradient observed in other species in which *oriC* is localized at one cell pole [3]. The absence of a ParB·*parS* nucleoprotein complex at the centromere (see below) might explain this non-polarized ParA_Bb_ pattern in AP *B. bacteriovorus* cells. Furthermore, our observations of ParA_Bb_ and ParB_Bb_ during chromosome replication and partitioning in the filamentous predator cell revealed several dynamic ParA_Bb_ “clouds” between segregating ParB_Bb_ foci, demonstrating that multiple ParABS systems are active simultaneously in *B. bacteriovorus*.

In contrast to binary-dividing species like *C. crescentus*, the partitioning complexes do not move across the predator cell from one pole to another. It is still unclear how the segregation of each chromosome is spatially confined within the filamenting cell body (which is filled with other replicating and segregating copies of the chromosome) to achieve the clear “beads-on-a-string” pattern of ParB foci ([48] and this study). We found that perturbations in the ParABS system, such as alterations of the ParA:ParB balance, prevent this exquisite arrangement of newly synthesized centromeres. In other bacteria, specific hub proteins at the cell poles polarize the choreography of ParABS-dependent segregation [29], but no homologs of these proteins were identified in *B. bacteriovorus*. Future research will determine whether this bacterium uses novel polar landmarks and other mechanisms to spatially organize chromosome partitioning along the mother cell.

### Coupling centromere organization to cell cycle progression

Our previous work uncovered a striking difference between the ParB protein in *B. bacteriovorus* and those characterized in other species. Whereas ParB homologs always localize at the *parS* site [35,37,56– 67], ParB_Bb_ is unable to accumulate at the chromosomal centromere in non-replicative cells, independently of its protein levels [48]. This study offers unambiguous support to this finding by showing that (i) the unique on-off localization pattern of ParB_Bb_ is independent of the fluorescent tag, and (ii) the protein fusions used to monitor ParB_Bb_ subcellular localization and abundance represent reliable reporters of the native protein during the entire *B. bacteriovorus* cell cycle. Furthermore, we identified ParB_Bb_ as a bona-fide CTPase like several other ParB homologs [10,13,74,78], which strongly suggests that ParB_Bb_ accumulation at the centromere results from its nucleation on *parS* and CTP-dependent sliding on adjacent DNA, while subsequent CTP hydrolysis unloads the protein.

Remarkably, we found that the absence of ParB_Bb_·*parS* complex depends on the chromosomal context of the centromere, as ParB_Bb_ formed a focus in *B. bacteriovorus* AP cells when *parS* was present on a plasmid. This result indicates that fluctuations in the cellular concentration of CTP, if any, are not responsible for the cell cycle-dependent localization of ParB_Bb_. It also hints that ParB_Bb_ has the capacity to bind and spread from *parS* in the absence of ongoing chromosome replication, even though the first ParB_Bb_ focus systematically appears after DNA replication initiation in GP cells [48]. Moreover, our data reveal that the direct chromosomal context of *parS* sites has a major role in preventing ParB_Bb_ complex formation, as ParB_Bb_ lost its ability to accumulate on the plasmidic *parS* when this sequence was flanked by short DNA stretches adjacent to *parS1* or *parS2* on the chromosome. This intriguing effect was only observed in *B. bacteriovorus,* and not in *E. coli* cells. Therefore, we propose that the local topology of the DNA in the immediate vicinity of the *parS* sequences, possibly induced by *B. bacteriovorus*-specific proteins during the GP-AP transition, might interfere with *parS* binding and/or the spreading of ParB_Bb_. Identifying the specific inhibition that prevents ParB_Bb_ from accumulating at the centromere in a cell-cycle and chromosomal context-dependent manner will certainly reveal additional complexity in bacterial centromere organization.

### Cell cycle-dependent fluctuations of the ParA and ParB protein balance

We found that ParB_Bb_ proteins accumulate during the growth of *B. bacteriovorus* but are nearly absent in newborn predator cells. Our data suggest that ParB_Bb_ is specifically degraded in a cell cycle-dependent manner. Although the co-expression of *parA_Bb_* and *parB_Bb_* is turned off during AP, protein concentration is not expected to change between the mother cell and the newborn cells immediately after cell division. However, ParB_Bb_ protein levels dropped within minutes at the GP-AP transition, while ParA_Bb_ levels remained stable within that time window. Whereas there has been no report of a mechanism involved in regulated ParB degradation in other bacteria, a recent study identified ParB as a candidate substrate of the conserved Lon protease in *C. crescentus* [79]. Future investigation will be needed to reveal the potential proteolytic mechanism that depletes ParB_Bb_ at the end of the *B. bacteriovorus* growth phase.

Nevertheless, this behavior appears to be crucial for *B. bacteriovorus* cell cycle progression, as higher levels of ParB_Bb_ led to cellular dysfunction, unless ParA_Bb_ was also overproduced. This observation suggests that *B. bacteriovorus* requires relatively higher levels of ParA_Bb_ than ParB_Bb_ in AP cells. Since additional, segregation-unrelated functions have been reported for ParA in other species (e.g., in transcription regulation) [80,81], it is tempting to speculate that ParA_Bb_ performs a ParB_Bb_-independent role during the G1 stage of the predator cell cycle. Finally, the compensatory effect of the double ParA_Bb_ and ParB_Bb_ overproduction may be related to the fine-tuned interplay of these proteins during the replicative phase, as excess ParA_Bb_ or ParB_Bb_ could impede proper pulling of chromosomal origins, resulting in the observed phenotypes. While studies in *Streptomyces coelicolor* have shown developmental stage-specific modulation of ParA and ParB protein levels resulting from the differential control of *parAB* transcription [42], to the best of our knowledge, our study is the first to report fluctuation of the ParA:ParB protein ratio across the lifecycle of a bacterium.

Altogether, our study sheds new light on the intricate dynamics and adaptation of the conserved ParABS system in bacteria. Our work hints that investigating the ParABS system in organisms with distinctive lifestyles will unveil new levels of complexity in centromere organization, chromosome segregation and cell cycle control.

## Materials and Methods

### Strains

The strains and plasmids used in this study are listed in Table S1, along with their respective construction methods (Table S2). Standard molecular cloning techniques were employed, and DNA assembly was conducted using the NEBuilder HiFi mix from New England Biolabs. All oligos used in the study can be found in Table S3. The *B. bacteriovorus* strains were produced from the wild-type HD100 strain, while the *E. coli* strains employed as prey were generated from MG1655. Microscopy, conjugation, and bacterial two-hybrid assays were conducted utilizing *E. coli* MG1655, S17-λpir, BL21, and BTH101. In addition, S17-λpir-derived *E. coli* strains were used for mating purposes. All plasmids were introduced into *B. bacteriovorus* through mating, following the methodology described below. Scarless allelic replacements into the HD100 chromosome were achieved via a two-step recombination approach using a pK18mobsacB-derived suicide vector. Chromosomal modifications were screened by PCR and verified by Sanger DNA sequencing, following the protocol detailed in [48].

### Routine culturing of B. bacteriovorus and E. coli

*E. coli* strains were grown in LB medium. For imaging experiments, overnight starter cultures from single colonies were diluted at least 1:500 in fresh medium until exponential phase. *B. bacteriovorus* strains were grown as described in DNB medium (Dilute Nutrient Broth, Becton, Dickinson, and Company) supplemented with 2 mM CaCl_2_ and 3 mM MgCl_2_ salts in the presence of *E. coli* prey at 30°C with continuous shaking [82]. When appropriate, antibiotic-resistant *E. coli* strains were used as prey for overnight culturing of the corresponding antibiotic-resistant *B. bacteriovorus*. Kanamycin or gentamycin was added in liquid and solid media at 50 µg/ml or 10 µg/ml, respectively.

### Plasmid conjugation by mating

Mating was carried out between the *E. coli* S17-λpir donor strain carrying the plasmid to be conjugated and the *B. bacteriovorus* receiver strain, following the protocol detailed in [48]. The exponentially growing *E. coli* donor strains were harvested and washed twice in DNB medium before resuspending in DNB salts at a 1:10 ratio of the initial volume. This donor suspension was combined with an equal volume of fresh overnight lysate of a receiver HD100 strain. The mating mixture was then incubated for a minimum of 4 hours at 30°C with shaking before being plated on a selective medium using the double-layer technique. Single plaques were isolated, and transconjugants were validated through microscopy (when applicable), PCR, and sequencing.

### RT-PCR

RNA samples were collected at different times during synchronous predation to track the expression of *parAB* (*bd3905-bd3906*) and *dnaK* (*bd1298*) throughout the predatory cycle of *B. bacteriovorus* on *E. coli* prey. A NucleoSpin RNA kit (MACHEREY-NAGEL) was used to isolate RNA, following the manufacturer’s instructions. An additional DNase treatment was performed for 1 hour at room temperature. RNA quality was assessed by measuring the 260/280 nm and 260/230 nm absorbance ratios. Reverse transcription and RT-PCR were carried out utilizing the QIAGEN OneStep RT-PCR kit with the following thermocycling parameters: 50°C for 30 minutes, 94°C for 15 minutes, followed by 30 cycles of 94°C for 1 minute, 50°C for 1 minute, and 72°C for 1 minute, with a final step of 72°C for 10 minutes.

### Bacterial-two hybrid assay (BACTH)

Interaction between protein pairs was assessed by the adenylate cyclase-based bacterial two-hybrid technique, as detailed in [83]. Briefly, the proteins of interest were merged with the isolated T18 and T25 catalytic domains of the *Bordetella pertussis* adenylate cyclase. The two plasmids producing the fusion proteins were introduced into the BTH101 reporter strain, and co-transformants were incubated on a selective medium overnight at 30°C. A single colony from each co-transformation was inoculated into 400 μl LB medium supplemented with ampicillin (200 µg/ml), kanamycin (50 µg/ml), and IPTG (0.5 mM). After incubation overnight at 30°C, 5 μl of each culture was placed onto LB plates supplemented with ampicillin, kanamycin, IPTG (using the abovementioned concentrations), and X-gal (40 ng/µl) and incubated at 30°C. An interaction assay with pKT25-Zip and pUT18C-Zip, two Zip protein domains, was included as a positive control. The experiments were conducted in triplicate, and representative results are presented.

### Live-cell imaging

*B. bacteriovorus* were grown overnight with the appropriate *E. coli* prey, and antibiotics were used when needed. Subsequently, they were grown on wild-type MG1655 for at least one generation without antibiotics before the beginning of the imaging experiment. For snapshots of fresh AP *B. bacteriovorus*, the cells were deposited on 1.2% agarose pads prepared with DNB-salt media. Regarding snapshots of *E. coli* strains, overnight cultures were diluted at least 1:500 and allowed to grow to the exponential phase before being deposited on 1.2% agarose pads prepared with PBS buffer supplemented with 0.2% glucose, 0.2% casamino acids, 1 µg/ml thiamine, 2 mM MgSO_4_, and 0.1 mM CaCl2). For the time-lapse imaging of synchronous predation cycles, MG1655 *E. coli* cells were grown in 2TYE medium to the exponential phase (OD_600_ = 0.4-0.6), harvested at 5000 x *g* at room temperature for 5 minutes, washed twice, and resuspended in DNB medium. *E. coli* and *B. bacteriovorus* were then combined in a 1:3 to 1:5 volume ratio to allow the majority of prey cells to be infected. In all synchronous predation imaging experiments, the prey-predator mixing step was indicated as time 0. The cells were either directly deposited on DNB-agarose pads for imaging or left shaking at 30°C before imaging for the indicated durations. In time-lapse experiments, the same fields of view on the pad were imaged at regular intervals, with the enclosure temperature set to 27°C. When appropriate, prior to imaging, cells were incubated for 5 minutes with DAPI (Life Technologies) at a final concentration of 5 µg/ml for nucleoid staining experiments.

### Image acquisition

Phase contrast and fluorescence images were obtained using a Nikon Ti2-E fully-motorized inverted epifluorescence microscope (Nikon) equipped with a CFI Plan Apochromat l DM 100x 1.45/0.13 mm Ph3 oil objective (Nikon), a Sola SEII FISH illuminator (Lumencor), a Prime95B camera (Photometrics), a temperature-controlled light-protected enclosure (Okolab), and filter-cubes for DAPI (32 mm, excitation 377/50, dichroic 409, emission 447/60; Nikon), mCherry (32 mm, excitation 562/40, dichroic 593, emission 640/75; Nikon), and GFP (32 mm, excitation 466/40, dichroic 495, emission 525/50; Nikon). Multi-dimensional image acquisition was supervised using the NIS-Ar software (Nikon). The pixel size was 0.074 µm using the 1.5X built-in zoom lens of the Ti2-E microscope. The same LED illumination and exposure times were applied when capturing images of various strains and/or conditions in one experiment and were set to a minimum for time-lapse acquisitions to limit phototoxicity.

### Image processing

To prepare figures, images were processed using FIJI [84], with contrast and brightness settings being kept consistent for all regions of interest in each figure unless stated otherwise. Denoising (Denoise.ai, Nikon) was applied to all phase contrast and fluorescence channels for **Fig 1B, 1E, and 1F** to improve the display of time-lapse images captured with low exposure (necessary to preserve cell viability). Figures were constructed and labeled using Adobe Illustrator. All outlines for the AP *B. bacteriovorus* were obtained with Oufti or drawn manually in Adobe Illustrator for bdelloplasts.

### Protein overexpression and purification

Expression and purification of ParB_Bb_ with a C-terminal His-tag were conducted as follows. *E. coli* BL21(DE3) cells containing pET21, which expresses C-terminal His-tagged ParB_Bb_,were cultivated at 37°C in autoinducing media supplemented with ampicillin (200 µg/ml). Following a 5-hour induction, cells were harvested by centrifugation and suspended in 35 mL lysis buffer (100 mM Tris-HCl [pH8], 300 mM NaCl, 5% glycerol (v/v), supplemented with a protease inhibitor cocktail (Complete, Roche)). The suspended cells were stored at 20 °C. The frozen cells were thawed on ice and lysed using three passages through a French pressure cell at 1500 psi. The suspension was then centrifuged for 15 minutes at 40 000 x *g* at 4 °C, and the supernatant was filtered through 0.45 µm filters before loading onto a 1 mL HisTrap column (GE Healthcare) pre-equilibrated with buffer A (100 mM Tris-HCl [pH8], 300 mM NaCl, and 5% [v/v] glycerol). The protein was eluted from the column using an increasing imidazole gradient (15–300 mM) in the same buffer. As a final purification step, size-exclusion chromatography was performed using a HiLoad 16/60 Superdex 75 column (GE Healthcare) with buffer B (100 mM Tris-HCl [pH8], 300 mM NaCl, and 5% [v/v] glycerol). The ParB_Bb_-containing fractions were pooled and analyzed for purity using SDS-PAGE. Glycerol was added to the ParB_Bb_ fractions to a final volume of 10 % (v/v), followed by 10 mM EDTA and 1 mM DTT. The purified ParB_Bb_ was aliquoted, snap-frozen in liquid nitrogen, and stored at –80 °C.

### Construction of DNA substrates for BLI assays

All DNA constructs were designed using VectorNTI (ThermoFisher) and synthesized chemically (gBlocks dsDNA fragments, IDT). To produce a linear biotinylated 40 bp DNA substrate, complementary oligos (with and without biotin) were heated at 98°C for 5 min before being left to cool down to room temperature overnight to form 50 mM double-stranded DNA. A 180 bp DNA construct was created with M13F and M13R homologous regions at each end. To produce a dual biotin-labeled DNA substrate, PCR reactions were carried out using a 2x GoTaq PCR master mix (Promega), biotin-labeled M13F and biotin-labeled M13R primers, and gBlocks fragments as a template. PCR products were separated by electrophoresis and then purified from the gel.

### Measurement of protein-DNA interaction by bio-layer interferometry (BLI)

Bio-layer interferometry experiments were performed using a BLItz system equipped with Dip-and-Read Streptavidin Biosensors (Molecular Devices) as previously described [15]. The streptavidin biosensor was first equilibrated in a low-salt binding buffer (100 mM Tris-HCl (pH 8), 100 mM NaCl, 1 mM MgCl2, and 0.005% Tween 20) for at least 10 min before each experiment. Biotinylated double-stranded DNA (dsDNA) was then immobilized onto the surface of the biosensor through a cycle of baseline (30 s), association (120 s), and dissociation (120 s). During the association phase, different concentrations of ParB_Bb_ dimers, with or without NTPs at varying concentrations, were added to the binding buffer and allowed to associate with the immobilized DNA for 120 s. Finally, the sensor was transferred into a protein-free binding buffer to monitor the dissociation kinetics for 120 s. The sensor was recycled by dipping in a high-salt buffer (100 mM Tris-HCl (pH 8), 1000 mM NaCl, 1 mM MgCl_2_, and 0.005% Tween 20) for 5 min to remove bound ParB_Bb_. All sensorgrams were recorded and analyzed using the BLItz analysis software (BLItz Pro version 1.2, Molecular Devices) and replotted using DataGraph for presentation. The data were fitted using a one-site-specific binding model in GraphPad Prism Version 5.04 to calculate the binding affinity constant Kd. All experiments were conducted at least in triplicate, and a representative sensorgram was presented in each figure.

### DNA preparation for EnzCheck phosphate assay

A 20 bp palindromic single-stranded DNA fragment (*parS_Bb_*: GGA<u>TGTTCCACGTGGAACA</u>TCC or *parS_Cc_*: GGA<u>TGTTTCACGTGAAACA</u>TCC; 100 mM in 1 mM Tris-HCl pH 8.0, 5 mM NaCl buffer) was heated at 98°C for 5 min before being left to cool down to room temperature overnight to form 50 µM double-stranded DNA. The *parS_Bb_* and *parS_Cc_* sequences are underlined.

### Measurement of NTPase activity by EnzCheck phosphate assay

The EnzCheck Phosphate Assay Kit from Thermo Fisher was used to measure NTP hydrolysis. A reaction buffer containing 1 mM NTP and 1 µM of ParB_Bb_ was analyzed in a Biotek EON plate reader at 25°C 15 hours with readings taken every minute. The reaction buffer consisted of 740 µL of ultrapure water, 50 µL of a 20x customized reaction buffer (100 mM Tris pH 8.0, 2 M NaCl, and 20 mM MgCl_2_), 200 µL of MESG substrate solution, and 10 µL of purine nucleoside phosphorylase (1 unit). Control reactions were performed with buffer only, buffer plus protein, or buffer plus NTP only. The plates were shaken continuously at 280 rpm for 15 hours at 25°C. Each assay was performed at least in triplicate. Data analysis was done using DataGraph, and the NTPase rates were calculated by fitting a linear regression in DataGraph.

### Western blot analysis

For Western blot analysis, the samples were prepared following the method used in [48], starting from 1.5 mL cleared predation lysates. Samples were collected at indicated time intervals for the time course analysis. NuPage Bis-Tris SDS precast polyacrylamide gels (Invitrogen) were used to load the samples and were run at 190 V for 50 minutes in the NuPAGE MOPS SDS running buffer. Standard Western blotting procedures were followed using primary polyclonal antibodies against ParB_Bb_ (rabbit sera obtained after injecting purified ParB_Bb_-6xhis protein; CER Group, Marloie, Belgium), GFP tag (JL-8 monoclonal antibody from Takara), and mCherry (polyclonal antibody from Thermo Fisher, product # PA5-34974). Detection of antibody binding was performed by visualizing chemiluminescence from the reaction of horseradish peroxidase with luminol, and chemiluminescence was imaged with an Image Quant LAS 500 camera (GE Healthcare). Secondary antibodies were goat anti-mouse IgG-peroxidase antibody (Sigma) for JL-8 and goat anti-rabbit IgG-peroxidase antibody (Sigma) for mCherry and ParB_Bb_. The images were analyzed using ImageJ, and the figures were assembled and annotated using Adobe Illustrator.

### Cell, nucleoid, and spot detection from microscopy images

The automated cell detection tool Oufti [85] was used to detect outlines of AP *B. bacteriovorus* cells, uninfected *E. coli* cells, or entire bdelloplasts with subpixel precision from phase contrast images. Fluorescent signals were added to cell meshes after background subtraction. Oufti was also used to detect diffraction-limited fluorescent foci and nucleoids with subpixel precision from fluorescence images, using the spotDetection and objectDetection modules. The detected spots and objects were added to the corresponding cell in the Oufti cell lists, including features related to coordinates, morphology, and intensity. The same optimized nucleoid detection parameters were used to ensure consistency and comparisons as done previously [48]. Parameters for spot detection were optimized for each dataset or control set of images.

### Quantitative image analysis from cell meshes

Fluorescence-related analysis, nucleoids, spots-related information, and other properties of individual cells based on microscopy images were extracted from Oufti cellLists data and plotted using custom codes in MATLAB (Mathworks), described below.

### Fluorescence profiles

The custom Matlab script MeanIntProfile.m was used to obtain mean relative fluorescence profiles. Briefly, the fluorescence profile of each cell (corresponding to the array of fluorescence intensity per cell segment provided by the relevant signal field in the Oufti cellList) was first normalized by the corresponding array of *steparea* values (corresponding to the area of each segment of the cell), then divided by their sum to obtain relative fluorescence values for each cell (to account for potential concentration differences between cells). When needed, arrays of relative fluorescence were oriented based on the position of the maximal fluorescence intensity of the indicated signal in each cell half. Cell length vectors were normalized from 0 to 1, and the corresponding relative fluorescence profiles from individual cells were interpolated to a fixed-dimension vector and concatenated before averaging.

### Fluorescence intensity analysis of time-lapse experiments

Bdelloplasts outlines were detected in the first frame of the time-lapse using Oufti and copied to all following frames using MATLAB. The resulting time-lapse cell list was reused in Oufti after background subtraction to add the fluorescence signal(s) to the bdelloplasts. The mean fluorescence profiles per cell over time were computed using a custom Matlab script, which extracts and plots the total fluorescence per cell (here bdelloplast) divided by the area of the corresponding cell (bdelloplast) over time.

### Kymographs and demographs

Demographs of relative fluorescence intensity in cells sorted by length were plotted as in [48,86]. When needed, arrays of relative fluorescence intensity values were oriented based on the position of the maximal fluorescence intensity of the indicated signal in each cell half. Kymographs were obtained using the built-in kymograph function in Oufti [85].

### Nucleoid size measurement

To measure nucleoid size, we used the objectDetection module in Oufti [85]. We considered the nucleoid area as a proxy for nucleoid size, based on previous work [87], which demonstrated that variations do not influence nucleoid area measurements in DAPI signal intensity. The nucleoid area was obtained from the nucleoid area field in the Oufti cell lists and exported to MATLAB for all cells with a single nucleoid. The nucleoid area values were then converted to µm^2^ and used to generate violin plots of nucleoid area distributions using R [88].

### Statistical analyses

The sample sizes and the number of repeats are included in the figure legends. Means, standard deviations, and coefficients of variation (CV) were calculated in MATLAB (Mathworks), R, or Microsoft Excel.

## Acknowledgements and funding sources

We thank Charles de Pierpont for excellent technical support, Dr. Kilian Dekoninck and Dr. Laurie Thouvenel for advice on protein purification, Dr. Michaël Deghelt for reviewing the draft, and all members of the Laloux lab for stimulating discussions and critical reading of the manuscript.

J.K. is a Research Fellow of the F.R.S.-FNRS, G.L. is a Research Associate of the F.R.S.-FNRS. J.K. was a recipient of travel and short-stay research grants from the European Molecular Biology Organization and the F.R.S.-FNRS. This work was supported by the European Commission (ERC Starting Grant PREDATOR #802331 to G.L.), the Fonds Jacques Moulaert (Fondation Louvain), the Lister institute fellowship, the Wellcome Trust Investigator grant (221776/Z/20/Z to T.B.K.L), and the Biotechnology and Biological Sciences Research Council grant-in-add (BBS/E/J/000PR9791 to the John Innes Centre).

## Author contributions

J.K. performed all experiments and analysis; ParB_Bb_ biochemical characterization experiments were conducted by J.K. in the laboratory of T.L; all authors analysed the data; J.K. prepared the figures; J.K. and G.L. drafted the manuscript; all authors revised the manuscript; G.L. supervised the work and acquired funding.

