## Supplemental Figures S1-S5 for "Cell cycle-dependent organization of a bacterial centromere through multi-layered regulation of the ParABS system"

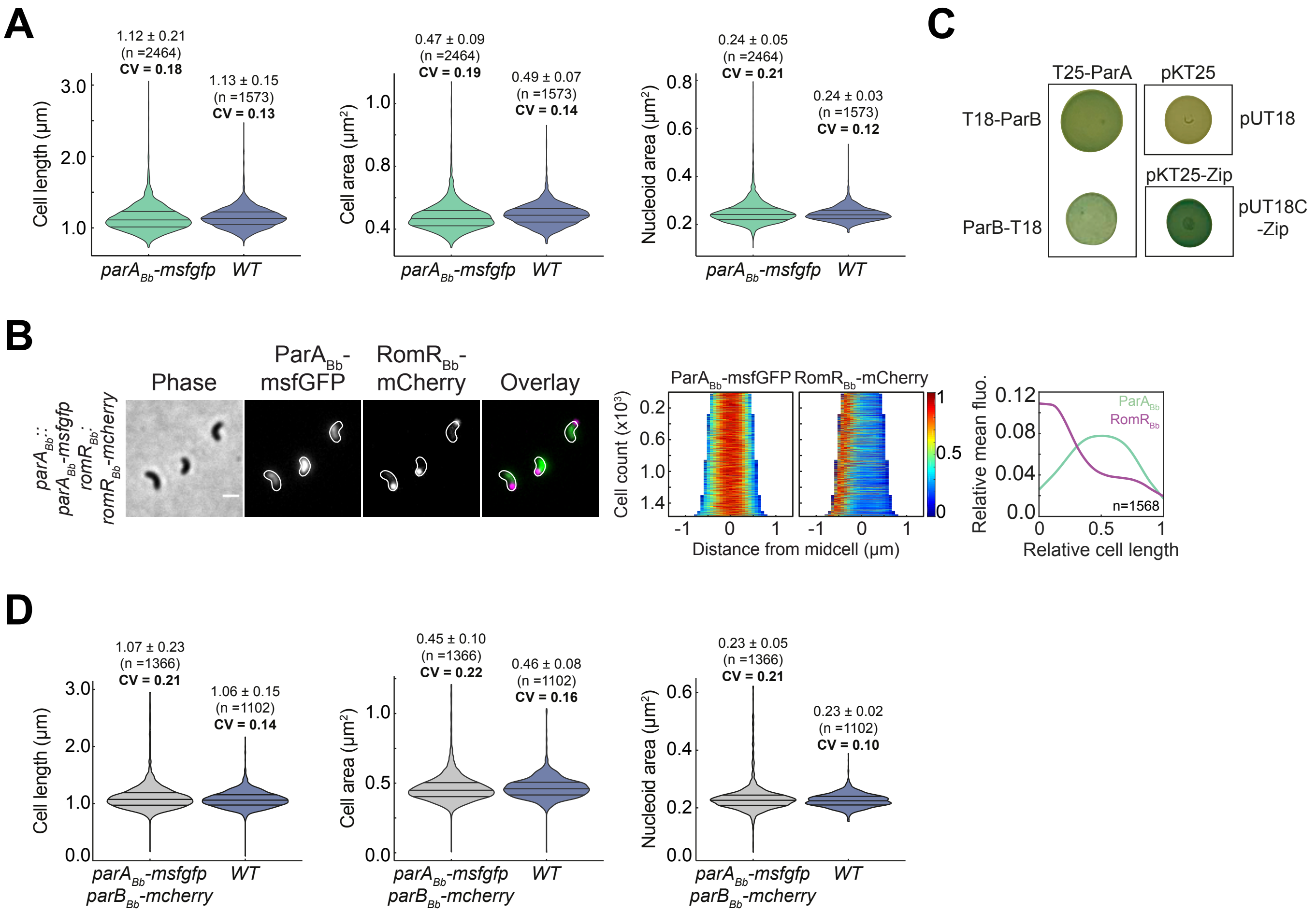

A
1.12 ± 0.21
(n =2464)
CV = 0.18
1.13 ± 0.15
(n =1573)
CV = 0.13
Cell length (µm)
parA$_{Bb}$-msfgfp
WT
0.47 ± 0.09
(n =2464)
CV = 0.19
0.49 ± 0.07
(n =1573)
CV = 0.14
Cell area (µm²)
0.24 ± 0.05
(n =2464)
CV = 0.21
0.24 ± 0.03
(n =1573)
CV = 0.12
Nucleoid area (µm²)
C
T25-ParA
pKT25
T18-ParB
pUT18
pKT25-Zip
ParB-T18
pUT18C
-Zip
B
Phase
ParA$_{Bb}$-msfGFP
RomR$_{Bb}$-mCherry
Overlay
parA$_{Bb}$::parA$_{Bb}$-msfgfp romR$_{Bb}$::romR$_{Bb}$-mcherry
ParA$_{Bb}$-msfGFP
RomR$_{Bb}$-mCherry
Cell count (x10³)
Distance from midcell (µm)
Relative mean fluo.
ParA$_{Bb}$
RomR$_{Bb}$
n=1568
Relative cell length
D
1.07 ± 0.23
(n =1366)
CV = 0.21
1.06 ± 0.15
(n =1102)
CV = 0.14
Cell length (µm)
parA$_{Bb}$-msfgfp
parB$_{Bb}$-mcherry
WT
0.45 ± 0.10
(n =1366)
CV = 0.22
0.46 ± 0.08
(n =1102)
CV = 0.16
Cell area (µm²)
0.23 ± 0.05
(n =1366)
CV = 0.21
0.23 ± 0.02
(n =1102)
CV = 0.10
Nucleoid area (µm²)


Figure S1

**Fig S1. ParA$_{Bb}$ subcellular localization and interaction with ParB$_{Bb}$. Related to Fig 1.**

(A) The endogenous ParA$_{Bb}$-msfGFP fusion is functional. Violin plots of cell length, cell area, and nucleoid area distributions in *parA$_{Bb}$::parA$_{Bb}$-msfgfp* (GL2134) and WT *B. bacteriovorus* strains, measured from cells in Fig 1E. The lines indicate the 25, 50, and 75 percent quantiles from bottom to top. Mean, standard deviation and coefficient of variation (CV) values are shown on top of the corresponding plot. n indicate the number of cells analyzed in a representative experiment.

(B) ParA$_{Bb}$ is not oriented towards any cell pole in AP cells. Left to right: representative phase contrast and fluorescence images of AP cells of *parA$_{Bb}$::parA$_{Bb}$-msfgfp romR::romR-mcherry* strain (GL2155); demographs of the corresponding fluorescent signals in the same cells sorted by length and oriented based on RomR-mCherry signal intensity; heatmaps represent relative fluorescence intensities; mean pole-to-pole profiles of relative fluorescence intensity of the corresponding fusions in the same cells. Scale bar is 1 µm.

(C) ParA$_{Bb}$ and ParB$_{Bb}$ interact in a bacterial two-hybrid assay. BTH101 reporter cells producing the indicated proteins fused to the T18 or T25 adenylate cyclase domain were spotted on X-gal agar plates supplemented with IPTG. Interaction between two proteins results in blue colony color. The Zip-Zip interaction serves as a positive control.

(D) A strain in which both native ParA$_{Bb}$ and ParB$_{Bb}$ are labeled displays morphology and chromosome aspect ratio similar to wild-type. Violin plots of cell length, cell area, and nucleoid area distributions in *parA$_{Bb}$::parA$_{Bb}$-msfgfp parB$_{Bb}$::parB$_{Bb}$-mcherry* (GL2154) and WT *B. bacteriovorus* strains, measured from cells in Fig 1G. The lines indicate the 25, 50, and 75 percent quantiles from bottom to top. Mean and standard deviation values are shown on top of the corresponding plot. n indicate the number of cells analyzed in a representative experiment. All experiments were performed at least twice.

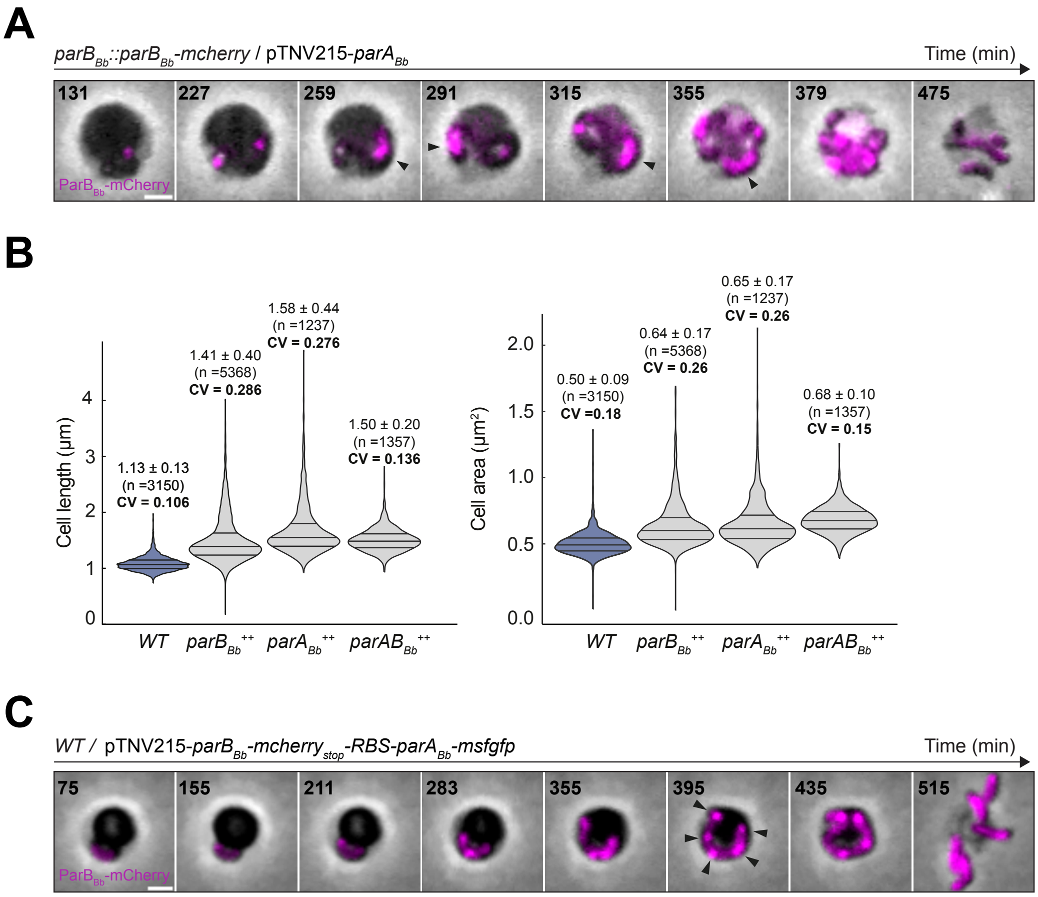
A
parBBb::parBBb-mcherry / pTNV215-parABb
Time (min)
131
227
259
291
315
355
379
475
ParBBb-mCherry
B
Cell length (µm)
1.13 ± 0.13
(n =3150)
CV = 0.106
1.41 ± 0.40
(n =5368)
CV = 0.286
1.58 ± 0.44
(n =1237)
CV = 0.276
1.50 ± 0.20
(n =1357)
CV = 0.136
Cell area (µm²)
0.50 ± 0.09
(n =3150)
CV =0.18
0.64 ± 0.17
(n =5368)
CV = 0.26
0.65 ± 0.17
(n =1237)
CV = 0.26
0.68 ± 0.10
(n =1357)
CV = 0.15
WT
parBBb++
parABb++
parABBb++
C
WT / pTNV215-parBBb-mcherrystop-RBS-parABb-msfgfp
Time (min)
75
155
211
283
355
395
435
515
ParBBb-mCherry

Figure S2

**Fig S2. Proper balancing of *parAB*$_{Bb}$ expression contributes to progressive *ori* segregation. Related to Fig 2.**

(A) Overexpression of *parA*$_{Bb}$ leads to chromosome segregation defects. *B. bacteriovorus* strain *parB*$_{Bb}$*::parB*$_{Bb}$*-mcherry* / pTNV215-*parA*$_{Bb}$ (GL2129) was mixed with prey and imaged in time-lapse after 120 min with 8-min intervals. Left: phase contrast and fluorescence images of selected time points; arrowhead points to an altered ParB$_{Bb}$-mCherry behavior (patches instead of well-separated foci) during the cell cycle.

(B) Overexpression of *parB*$_{Bb}$ or *parA*$_{Bb}$ leads to pronounced phenotypes, largely rescued by overexpression of both. Violin plots of cell length, cell area, and nucleoid area distributions in WT / pTNV215-*parB*$_{Bb}$ (*parB*$_{Bb}^{++}$, GL1261); WT / pTNV215-*parA*$_{Bb}$ (*parA*$_{Bb}^{++}$, GL1460); WT / pTNV215-*parB*$_{Bb}$-*parA*$_{Bb}$ (*parAB*$_{Bb}^{++}$, GL1004) and WT *B. bacteriovorus* from cells in Fig 2B. The lines indicate the 25, 50, and 75 percent quantiles from bottom to top. Mean, standard deviation and coefficient of variation (CV) values are shown on top of the corresponding plot. n indicate the number of cells analyzed in a representative experiment.

(C) Overexpression of both *parA*$_{Bb}$ and *parB*$_{Bb}$ has no obvious effect on *ori* segregation. *B. bacteriovorus* strain WT / pTNV215-*parB*$_{Bb}$*-mcherry*$_{stop}$*-RBS-parA*$_{Bb}$*-msfgfp* (GL1004) was mixed with prey and imaged in time-lapse after 75 min with 8-min intervals. Left: phase contrast and fluorescence images of selected time points; arrowheads point to well-distributed ParB$_{Bb}$-mCherry foci (which mark chromosomal *ori*) during the cell cycle; ParA$_{Bb}$-msfGFP not shown for simplicity. All experiments were performed at least twice. Scale bars are 1 µm.

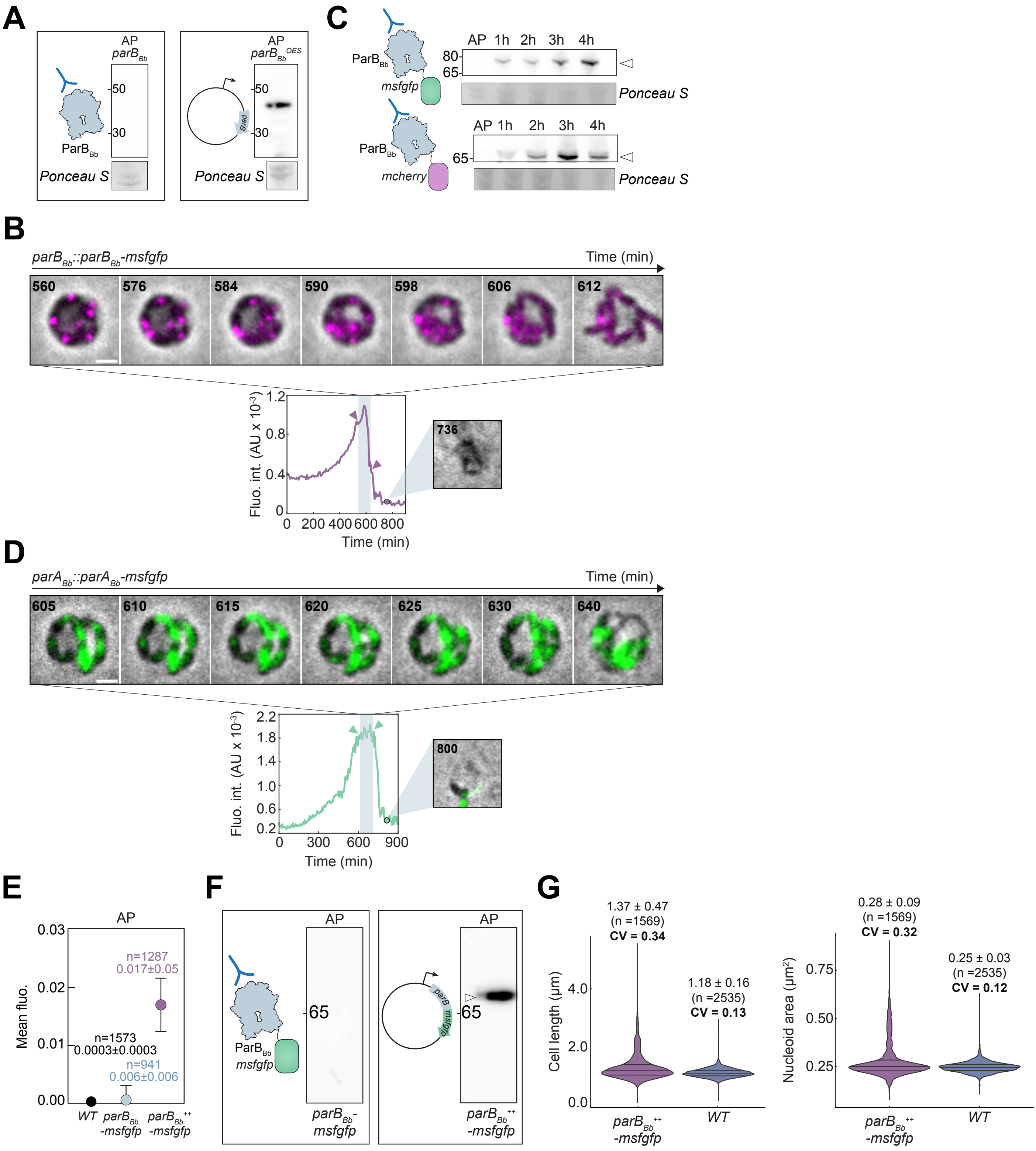
A
AP
$parB_{Bb}$
50
30
$ParB_{Bb}$
Ponceau S
AP
$parB_{Bb}^{OES}$
50
30
Ponceau S
C
AP 1h 2h 3h 4h
80
65
$ParB_{Bb}$
msfgfp
Ponceau S
AP 1h 2h 3h 4h
65
$ParB_{Bb}$
mcherry
Ponceau S
B
$parB_{Bb}$::$parB_{Bb}$-msfgfp
Time (min)
560
576
584
590
598
606
612
Fluo. int. (AU x $10^{-3}$)
Time (min)
736
D
$parA_{Bb}$::$parA_{Bb}$-msfgfp
Time (min)
605
610
615
620
625
630
640
Fluo. int. (AU x $10^{-3}$)
Time (min)
800
E
AP
Mean fluo.
n=1287
0.017±0.05
n=1573
0.0003±0.0003
n=941
0.006±0.006
WT
$parB_{Bb}$-msfgfp
$parB_{Bb}^{++}$-msfgfp
F
AP
65
$ParB_{Bb}$
msfgfp
$parB_{Bb}$-msfgfp
AP
65
$parB_{Bb}^{++}$-msfgfp
G
1.37 ± 0.47
(n =1569)
CV = 0.34
1.18 ± 0.16
(n =2535)
CV = 0.13
Cell length (µm)
$parB_{Bb}^{++}$-msfgfp
WT
0.28 ± 0.09
(n =1569)
CV = 0.32
0.25 ± 0.03
(n =2535)
CV = 0.12
Nucleoid area (µm²)
$parB_{Bb}^{++}$-msfgfp
WT

Figure S3

**Fig S3. The levels of ParB$_{Bb}$ and ParA$_{Bb}$ vary differently during the cell cycle, and ParB$_{Bb}$ is unable to form foci when overproduced. Related to Fig 3.**

(A) Anti-ParB$_{Bb}$ antibody detects overproduced untagged ParB$_{Bb}$ in AP (right) - control experiment for Fig 3A and S3A (left). Western blots of whole-cell protein extracts from AP cells of *WT B. bacteriovorus* and a strain constitutively producing the untagged ParB$_{Bb}$ (GL1261) were probed with an anti-ParB$_{Bb}$ antibody represented in the schematics on the left.

(B) Same as in Fig 3B for the strain natively producing ParB$_{Bb}$-msfGFP (GL1654).

(C) The protein levels of natively produced ParB$_{Bb}$-msfGFP or ParB$_{Bb}$-mCherry detected with the same anti-ParB$_{Bb}$ antibody compare with the endogenous untagged ParB$_{Bb}$ profile during a synchronized *B. bacteriovorus* cell cycle. Western blots of whole-cell protein extracts from *B. bacteriovorus* strains *parB$_{Bb}$::parB$_{Bb}$-msfgfp* (GL1654) and *parB$_{Bb}$::parB$_{Bb}$-mcherry* (GL906) were probed with an anti-ParB$_{Bb}$ antibody, as represented in the schematics. Protein samples are isolated at time points throughout the predatory cell cycle: AP attack phase, 1h, 2h, 3h, and 4h after mixing with prey. Arrowhead indicates detected ParB$_{Bb}$ protein during the growth phase. Ponceau staining of the same membranes (where bands were most visible, ~30-50 kDa) is illustrated below each blot as a loading control. Molecular weight markers (kDa) are shown on the side.

(D) Same as in Fig 3B for the strain natively producing ParA$_{Bb}$-msfGFP (GL2134).

(E) Mean msfGFP fluorescence measured for AP cells of WT, *parB$_{Bb}$::parB$_{Bb}$-msfgfp* (*parB$_{Bb}$-msfgfp,* GL1654) and *WT* / pTNV215-*parB$_{Bb}$-msfgfp* (*parB$_{Bb}$-msfgfp$^{++}$,* GL1003). n indicate the number of cells analyzed in a representative experiment; mean values are represented. Error bars indicate standard deviations.

(F) Western blots of whole-cell protein extracts from AP cells of WT *B. bacteriovorus parB$_{Bb}$::parB$_{Bb}$-msfgfp* (*parB$_{Bb}$-msfgfp,* GL1654, left) and *WT* / pTNV215-*parB$_{Bb}$-msfgfp* (*parB$_{Bb}$-msfgfp$^{++}$,* GL1003, right) strains were probed with an anti-ParB$_{Bb}$ antibody represented in the schematics on the left. Arrowhead points to the ParB$_{Bb}$-msfGFP protein detected in AP when overproduced.

(G) Overproduction of ParB$_{Bb}$-msfGFP leads to morphological and chromosome segregation defects. Violin plots of cell length and nucleoid area distributions for the cells in Fig 3D and WT *B. bacteriovorus*. The lines indicate the 25, 50, and 75 percent quantiles from bottom to top. Mean, standard deviation and coefficient of variation (CV) values are shown on top of the corresponding plot. n indicate the number of cells analyzed in a representative experiment. All experiments were performed at least twice.

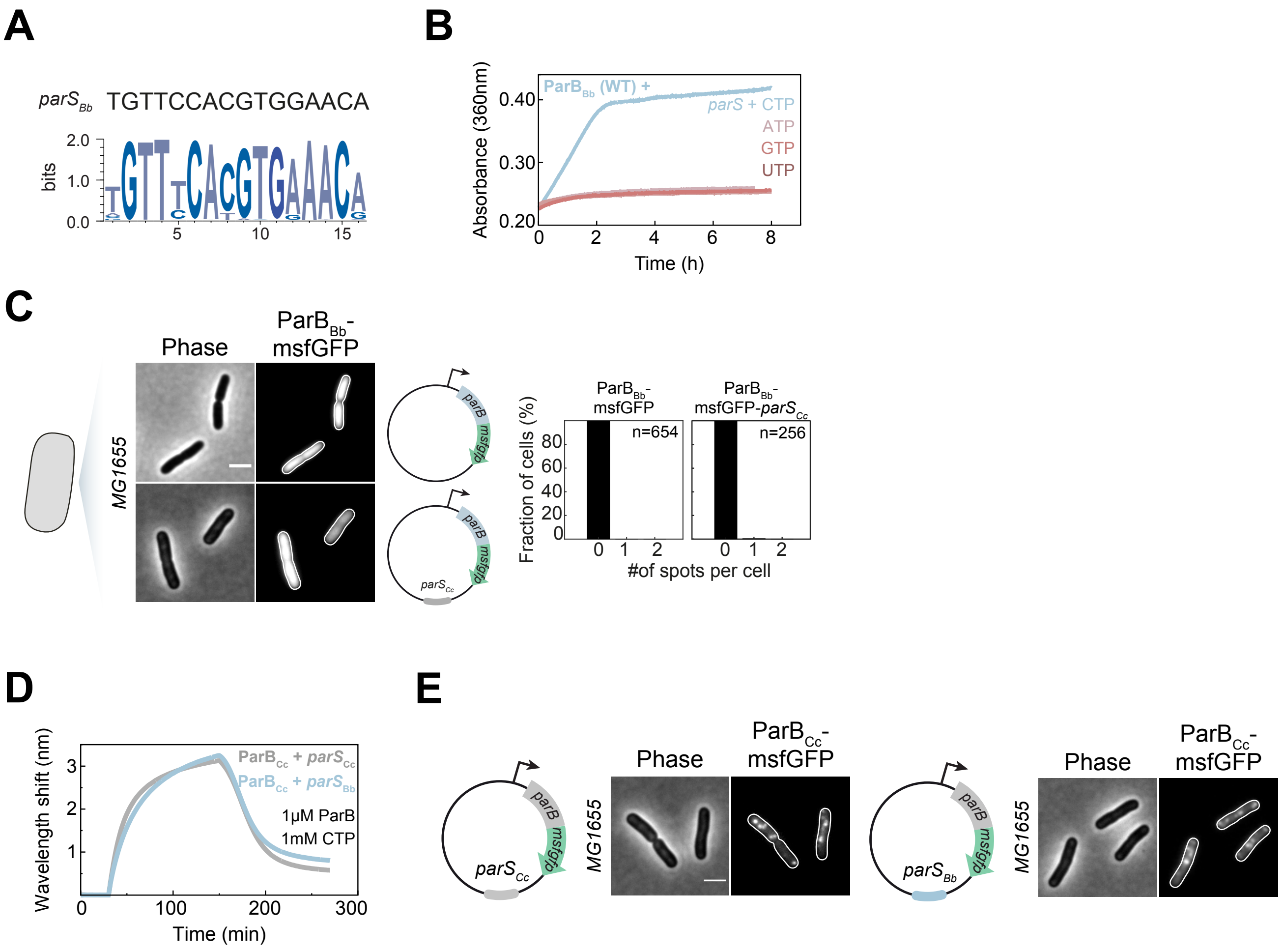
A
parS Bb TGTTCCACGTGGAACA
bits
2.0
1.0
0.0
5
10
15
B
Absorbance (360nm)
0.40
0.30
0.20
ParB Bb (WT) +
parS + CTP
ATP
GTP
UTP
0
2
4
6
8
Time (h)
C
Phase
ParB Bb-msfGFP
MG1655
parB
msfgfp
parS Cc
ParB Bb-msfGFP
ParB Bb-msfGFP-parS Cc
n=654
n=256
Fraction of cells (%)
80
60
40
20
0
0 1 2
#of spots per cell
D
Wavelength shift (nm)
ParB Cc + parS Cc
ParB Cc + parS Bb
1μM ParB
1mM CTP
0
100
200
300
Time (min)
E
Phase
ParB Cc-msfGFP
MG1655
parB
msfgfp
parS Cc
parS Bb

Figure S4

**Fig S4. Investigation of the $ParB_{Bb}$ features near the CTP binding pocket and their impact on *$parS_{Bb}$* recognition. Related to Fig 4.**

A) *$parS_{Bb}$* sequence matches with the consensus. Top: *parS* sequence from *B. bacteriovorus.* Bottom: a sequence logo generated by WebLogo 3.0 (10), using *parS* sequence alignments from (11) as input.

(B) $ParB_{Bb}$ is a CTPase. Continuous monitoring of the released inorganic phosphate (Pi) by recording the absorbance at 360 nm overtime at 25℃. The NTP hydrolysis of $ParB_{Bb}$ was also monitored in the presence of ATP, GTP, or UTP, with a 22 bp *$parS_{Bb}$* DNA duplex.

(C) $ParB_{Bb}$ does not cluster without *parS* or on non-cognate *parS.* Left: representative phase contrast and fluorescence images of MG1655 *E. coli* strain constitutively producing $ParB_{Bb}$-msfGFP from a plasmid carrying no *parS* sequence (GL1661) or the *C. crescentus parS* (*$parS_{Cc}$*; GL2024). Right: histograms representing the percentage of cells with zero, one, or two $ParB_{Bb}$-msfGFP foci in the same strains. The scale bar is 2 µm; schematics illustrate *$parB_{Bb}$-msfgfp* expression plasmids.

(D) ParB from *C. crescentus* ($ParB_{Cc}$) is more promiscuous to *parS* binding than $ParB_{Bb}$. BLI analysis of the interaction between 1 µM $ParB_{Cc}$ and a 40 bp cognate *$parS_{Cc}$* (grey) or a non-cognate *$parS_{Bb}$* (blue). $ParB_{Cc}$ binds both *parS* sequences.

(E) $ParB_{Cc}$ can bind to its cognate *$parS_{Cc}$* and a non-cognate *$parS_{Bb}$* in *E. coli*. Left: representative phase contrast and fluorescence images of MG1655 *E. coli* strain constitutively producing $ParB_{Cc}$-msfGFP from a plasmid carrying *$parS_{Cc}$* (left, GL2025) or *$parS_{Bb}$* (right, GL2026); fluorescent foci are observed in both cases. The scale bar is 2 µm; schematics illustrate the *$parB_{Bb}$-msfgfp* expression plasmid.

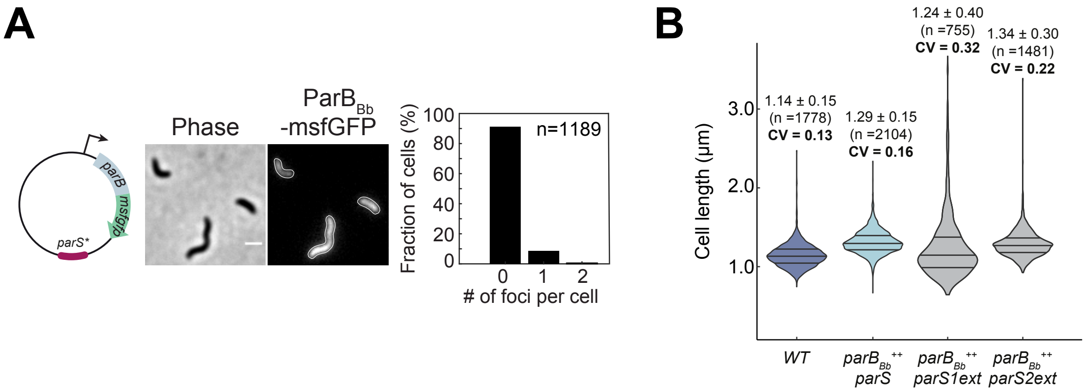
A
Phase
ParBBb
-msfGFP
parB
msfgfp
parS*
Fraction of cells (%)
n=1189
100
80
60
40
20
0
0
1
2
### of foci per cell
B
Cell length (µm)
3.0
2.0
1.0
1.14 ± 0.15
(n =1778)
CV = 0.13
1.29 ± 0.15
(n =2104)
CV = 0.16
1.24 ± 0.40
(n =755)
CV = 0.32
1.34 ± 0.30
(n =1481)
CV = 0.22
WT
parBBb++ parS
parBBb++ parS1ext
parBBb++ parS2ext

Figure S5

**Fig S5. Nucleation of ParB$_{Bb}$ on plasmidic parS$_{Bb}$ is specific; titration of the free ParB$_{Bb}$ by parS$_{Bb}$ rescues the *parB* overexpression phenotype. Related to Fig 5.**

(A) From left to right: representative phase contrast and fluorescence images of AP cells of WT *B. bacteriovorus* constitutively producing ParB$_{Bb}$-msfGFP from a plasmid carrying a mutated *–parS$_{Bb}$* (*parS$_{Bb}$*,* GL1541). Right: histogram representing the percentage of cells with zero, one, or two ParB$_{Bb}$-msfGFP foci in the same strain.

(B) Only the cytosolic excess ParB$_{Bb}$ is toxic for *B. bacteriovorus* AP cells. Violin plots of cell length for the cells in Fig 5A-C and WT *B. bacteriovorus*. The lines indicate the 25, 50, and 75 percent quantiles from bottom to top. Mean, standard deviation and coefficient of variation (CV) values are shown on top of the corresponding plot. n indicate the number of cells analyzed in a representative experiment; all experiments were performed at least twice.
